# Constitutive activation of canonical Wnt signaling disrupts choroid plexus epithelial fate

**DOI:** 10.1101/2020.12.08.415588

**Authors:** Arpan Parichha, Varun Suresh, Mallika Chatterjee, Aditya Kshirsagar, Lihi Ben-Reuven, Tsviya Olender, M. Mark Taketo, Velena Radosevic, Mihaela Bobic-Rasonja, Sara Trnski, Michael J. Holtzman, Nataša Jovanov Milošević, Orly Reiner, Shubha Tole

## Abstract

The choroid plexus (ChP) secretes cerebrospinal fluid and is critical for the development and function of the brain. In the telencephalon, the ChP epithelium (ChPe) arises from the *Wnt*-expressing cortical hem. Embryonic mouse and human ChPe both express nuclear β-CATENIN, a canonical Wnt signaling pathway effector, indicating that this pathway is active during ChPe development. Point mutations in human *β-CATENIN* result in the constitutive activation of canonical Wnt signaling. In a mouse model that recapitulates this perturbation, we report a loss of ChPe identity and an apparent transformation of the ChPe to a neuronal identity. Aspects of this phenomenon are recapitulated in human embryonic stem cell (hESC)-derived organoids. The ChPe is also disrupted when *β-Catenin* is conditionally inactivated in the mouse. Together, our results indicate that canonical Wnt signaling is required in a precise and regulated manner for normal ChPe development in the mammalian brain.

## Introduction

The Choroid plexus (ChP) produces cerebrospinal fluid (CSF) which plays an important role in the developing and adult brain. The CSF provides protective functions such as cushioning against mechanical damage, nutritive function, and signaling functions, providing molecular cues for the developing brain^1,2^. The ChP separates the blood from the CSF creating a checkpoint for regulating the entry and exit of factors^2,3^. The ChP is also the gateway for immune cells entering or exiting the brain^4^.

The ChP epithelium (ChPe) arises from adjacent neuroepithelial tissue in the telencephalon, diencephalon, and hindbrain. In the telencephalon, the non-neuronal ChPe and a sub-population of Cajal-Retzius (CR) neurons both arise from the cortical hem^5–7^, a midline telencephalic signaling center enriched in genes that belong to the BMP and WNT protein families. The telencephalic hem has been identified in several vertebrate classes^8^. In the opossum *Monodelphis domestica,* BrdU positive cells arise from the hem and populate the growing telencephalic ChPe in a “conveyor belt” like manner^9^. Consistent with this model, when the size of the cortical hem increases^10^, decreases^11^ or is missing altogether^12^, there is a parallel change in the extent of the ChP.

The mechanisms that regulate the specification of the ChPe are important from developmental, disease, and evolutionary perspectives. Obvious candidates for the control of ChPe development are the signaling molecules expressed in the neuroepithelial progenitor domains. The hem expresses several members of the Bmp and Wnt families^12,13^. When the expression of these signaling molecules in the hem is compromised, such as in the Gli3 null (Xt^J^) mutant, neither the ChP nor the hippocampus is specified normally^12,14^. Bmp signaling is critical for ChPe specification. The ChP is missing in the *BmpR1a* null mutant mouse^15,16^, and *Bmp4* is sufficient for ChPe induction *in vitro*^16,17^. The role of hem-derived Wnt ligands in ChPe development appears to be equally important, though not well explored. Wnt5a, expressed in the hem, appears to be required for maintaining the size and the cytoarchitecture of the ChPe^18^. A role for canonical Wnt signaling in ChPe development is suggested by the observation that the choroid plexus appears to be reduced in the *Wnt3a* null mutant^19^. Furthermore, the addition of WNT3a augments the ChPe-inducing ability of BMP4 in a three-dimensional aggregation culture derived from embryonic stem cells (ESCs)^17^.

In this study, we demonstrate that molecules downstream of canonical Wnt signaling, such as nuclear β-CATENIN, are present *in vivo* at embryonic stages in the human and mouse ChPe. Mutations in particular domains of human *β-CATENIN* have been reported to cause constitutive activation of this pathway since they interfere with β-CATENIN phosphorylation and consequent degradation^20^. Specifically, exon 3 of the *β-Catenin (Ctnnb1)* gene is a hotspot for several mutations, including point mutations and deletions associated with various forms of cancer^21^. The effects of these mutations on ChP-related disorders have not been studied, even though ChP papillomas have been reported to display upregulation of canonical Wnt signaling^22^. Therefore, we focused our studies on examining the effects of constitutive activation of β-CATENIN in the mouse ChP using a strategy in which exon 3 can be conditionally deleted. We discovered that this perturbation causes a loss of ChPe identity and, surprisingly, a gain of neuronal identity, together with comprehensive transcriptomic changes that are consistent with the activation of canonical Wnt signaling. We paralleled these studies in a human ESC (hESC)-derived organoid model in which treatment with a low level of a canonical Wnt agonist together with BMP4 resulted in a ChPe-like fate, but over-activation with high levels of the Wnt agonist recapitulated the loss of ChPe identity seen in the mouse. Loss of *β-catenin* in the mouse resulting in an absence of canonical Wnt signaling^23^ also disrupted ChPe development.

Together, our results indicate that controlled levels of canonical Wnt signaling is critical for ChPe specification and morphogenesis and that this mechanism appears to be conserved in humans and rodents.

## Results

### The human fetal ChPe and the embryonic mouse ChPe display components of canonical Wnt signaling

We examined whether the molecular machinery required for processing canonical Wnt signaling is present in the developing ChPe at the relevant stages in both human and mouse embryos. In humans, the telencephalic ChP develops between gestational weeks 8 to 12. We collected human fetal samples from GW11 and GW13 and performed immunohistochemical analysis for Wnt signaling components. The human ChP is identified by its fan-like morphology within the telencephalic ventricles (Figure 1A, B), and the presence of AQP1 and OTX2 that are established ChPe markers (Figure 1A-G). The canonical Wnt receptor, FZD1, is detected throughout the ChPe (Figure 1B). LEF1, AXIN2, and β-CATENIN are also present, consistent with canonical Wnt signaling. In particular, β-CATENIN was detected in the nucleus, (Figure 1D, E) indicating that canonical Wnt signaling is likely to be active in the human ChPe at early stages during ChP development.

**Figure 1.**
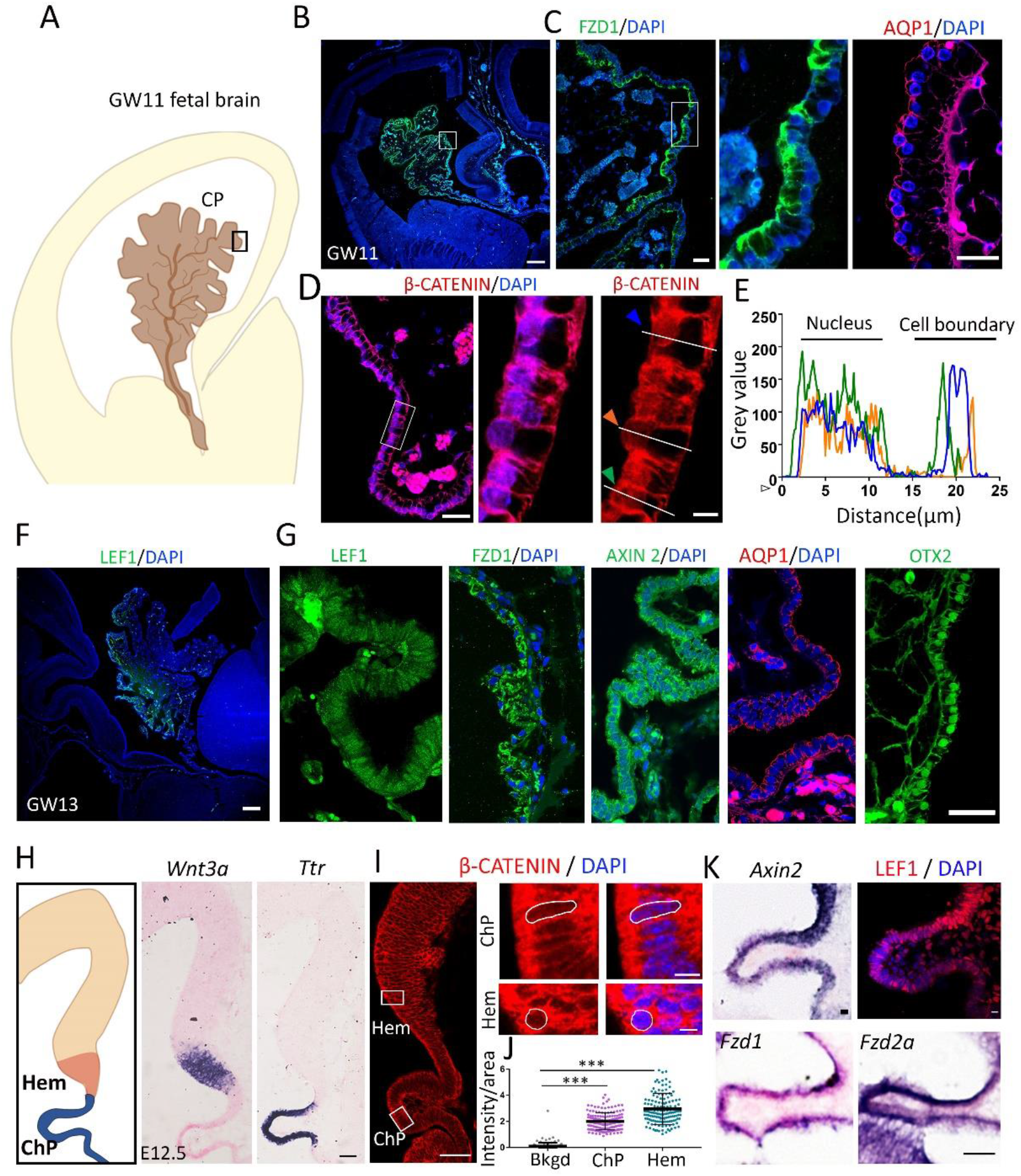
Canonical Wnt signaling components are present in the developing human and mouse choroid plexus epithelium. (A-G) Canonical Wnt signaling pathway molecules FZD1, β-CATENIN, LEF1, and AXIN2, and also ChPe markers OTX2 and AQP1 are detected at gestational week (GW) 11 (B-E) and GW13 (F and G) in the human choroid plexus epithelium. (D, E) β-CATENIN immunohistochemistry intensity measurements across three ChPe cells (identified by colored arrowheads in D corresponding to traces in E). β-CATENIN intensity is seen in the nucleus, indicative of active canonical Wnt signaling and the boundaries between cells. (H) In the E12.5 mouse telencephalon, *Wnt3a* and *Ttr* expression identifies the hem the ChPe, respectively. (I, J) Nuclear localization of β-CATENIN and its quantitation from 150 ChPe and hem nuclei (Bkgd, background; N=3). (K) Canonical Wnt signaling components *Axin2, Fzd1, Fzd2a*, and LEF1 are expressed in the E12.5 mouse ChPe. Statistical test (J): One Way ANOVA followed by post hoc Dunnett’s multiple comparison test * p < 0.05, ** p < 0.01, *** p < 0.001, ns if p-value> 0.05. Scale bars: 500 μm (B and F); 50 μm (D and I low mag, G, H, and K); 10 μm (D and I high mag).

In the embryonic day E12.5 mouse, the hem and the ChPe are identified by the expression of *Wnt3a* and the thyroid hormone T4 carrier transthyretin (*Ttr*), respectively (Figure 1H). The expression of several canonical Wnt signaling pathway molecules was detected in the ChPe including receptors such as *Frizzled 1* and *2a* (*Fzd1, Fzd2a*), a target gene *Axin2*, and the transcriptional regulators β-CATENIN and LEF1 were present in nuclei of ChPe cells (Figure 1I-K). These data suggest that the embryonic human and mouse ChPe have the capability of processing canonical Wnt signaling.

### Disrupting β-CATENIN function in the hem and ChPe

*Lmx1a* expression is specific to the hem and the ChPe within the telencephalon^24^ . To genetically manipulate the developing ChPe we used the Lmx1aCre line, which faithfully labels the hem lineage that is composed of Cajal-Retzius cells and ChPe cells, when crossed to the Ai9 reporter line^25^ (Figure 2A). Therefore, the Lmx1aCre line offers the opportunity of disrupting genetic mechanisms not only in the ChPe, but also in its domain of origin, the hem.

**Figure 2:**
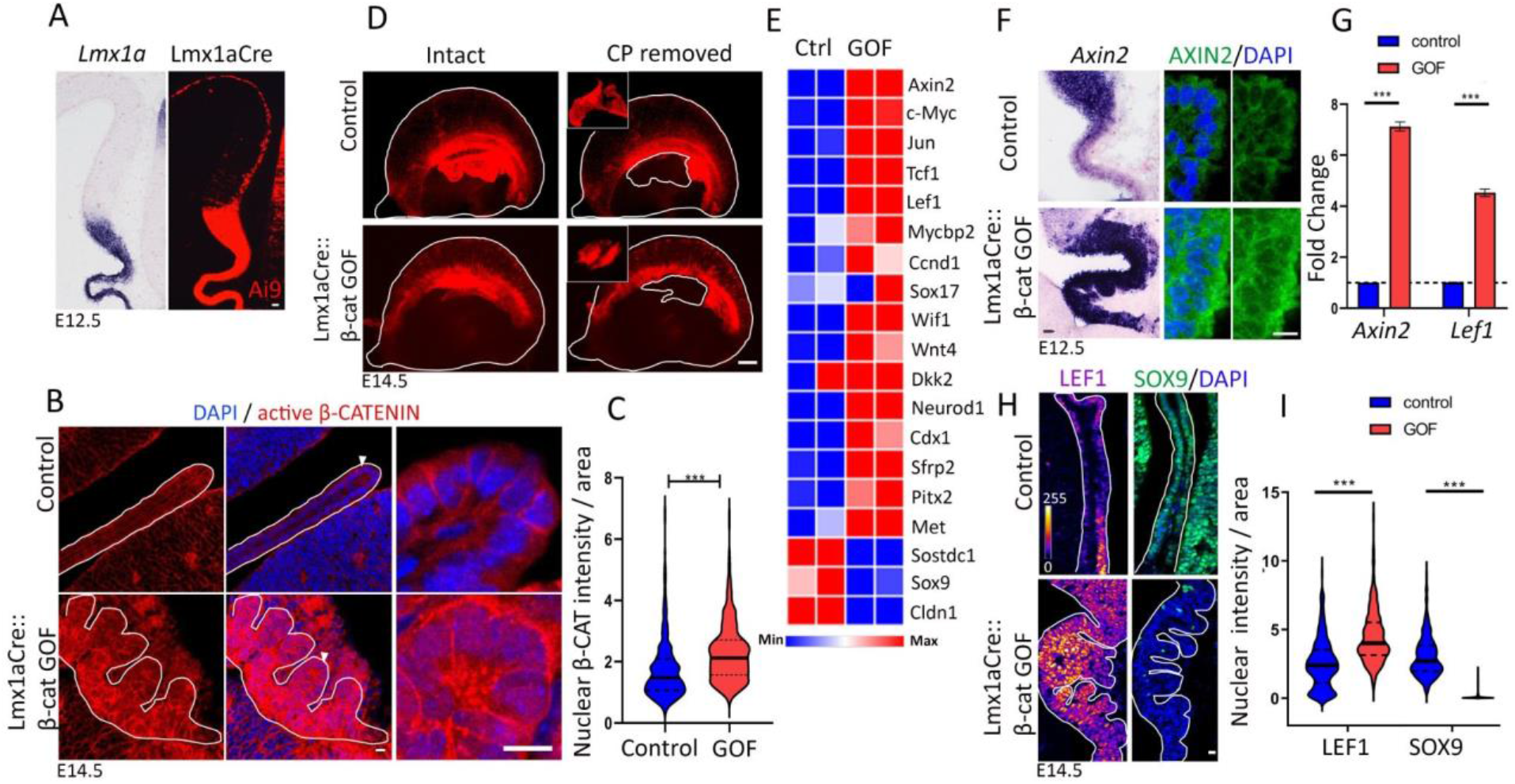
Stabilization of β-CATENIN leads to activation of canonical Wnt signaling. (A) At E12.5, in situ hybridization for *Lmx1a* shows endogenous expression; the Ai9 reporter labels the Lmx1a-expressing lineage, including the hem, ChPe, and hem-derived Cajal-Retzius cells. (B, C) At E14.5, nuclear localization of active (non-phosphorylated) β-CATENIN is increased in the Lmx1aCre:: β-cat GOF ChPe compared with controls as seen by (B) immunohistochemistry and (C) a violin plot representing quantitation of 294 nuclei (control) and 301 (GOF); N=3 for each genotype. (D) Microdissection of E14.5 Ai9-labeled telencephalic hemispheres to isolate ChP (inset) for RNA extraction. (E) Heat map of Wnt-signaling target genes from RNA-seq data from control and Lmx1aCre::β-Catenin GOF ChP. (F-I) In situ hybridization (Axin2), qPCR Axin2 and Lef1, and immunohistochemistry (AXIN2, LEF1, and SOX9) reveal up regulation of AXIN2 and LEF1 and downregulation of SOX9 in the Lmx1aCre:: β-cat GOF ChPe compared with controls. Bar graphs (G) represent mean ± SEM (N=3). (I) Violin plots quantifying the nuclear intensity of LEF1 and SOX9 (LEF1: N=5, 505 nuclei from control and 500 nuclei from Lmx1aCre:: β-cat GOF ChPe. SOX9 N=4, 457 nuclei from control and 444 nuclei from Lmx1aCre:: β-cat GOF ChPe). Statistical tests (C,G,I): Two-tailed unpaired multiple Student’s t-test with unequal variance; * p < 0.05, ** p < 0.01, *** p < 0.001, ns if p-value> 0.05. Scale bars: 10 μm (A, B, F & H); 100 μm (D).

We used Lmx1aCre together with two well-described mouse lines to introduce either gain-of-function (GOF) or loss-of-function (LOF) genetic perturbations in the *β-catenin* (*Ctnnb1*) gene. In *β-catenin* GOF mice, exon 3 of *β-catenin* is flanked by loxP sites, and Cre-mediated recombination results in deletion of a domain that contains phosphorylation sites for GSK3*β*, which would normally tag β-CATENIN for degradation. The recombined *β-catenin* allele encodes a protein that remains constitutively active^26^ (gain of function). This results in increased accumulation of active β-CATENIN within the nucleus of ChPe cells consistent with its role as a transcription factor^27^. Intensity quantification of the immunofluorescence within the nuclei of ChPe cells revealed that GOF mice have higher levels of active nuclear β-CATENIN than controls (Figure 2B, C).

We examined whether these perturbations resulted in transcriptional changes that are typical of canonical Wnt pathway activity. We micro dissected the ChP from E14.5 control and β-catenin GOF embryos, using the expression of the Ai9 reporter (Figure 2D). This gave us a relatively pure ChP preparation that was processed for bulk RNA-seq. 2439 genes were downregulated and 2163 genes upregulated in the ChP upon constitutive activation of β-CATENIN, using a 1.5-fold cutoff (p.adj < 0.05; Supplementary Table 1). KEGG pathway analysis identified the Wnt pathway as one of the major pathways dysregulated in the Lmx1aCre::β-catenin GOF ChP (Supplementary Figure S1). Consistent with this, multiple canonical Wnt pathway targets were positively regulated (*Ccnd1, Cdx1, Cldn1, c-Myc, Dkk2, Jun, Lef1, Met, Mycbp2, Neurod1, Pitx2, Sfrp2, Sox17, Tcf1, Wif1, Wnt4)*, and others were negatively regulated (*Sostdc1, Sox9,*) (Figure 2E; Supplementary Figure S1;^28,29^, https://web.stanford.edu/group/nusselab/cgi-bin/wnt/target_genes). We further examined well-established positively regulated targets of this pathway, *Axin2*, LEF1, and a negatively regulated target SOX9, by *in situ* hybridization, qPCR, and/or immunohistochemistry, in the control and Lmx1aCre::β-catenin GOF ChPe. Both mRNA and protein for *Axin2* and *Lef1* showed a marked increase in the GOF ChPe compared with controls. In contrast, SOX9, which is normally present in the ChPe, is greatly reduced in the β-catenin GOF ChPe, consistent with its suppression by canonical Wnt signaling^30^ (Figure 2 F-I).

We also examined the loss of function of β-CATENIN using Lmx1aCre using a well-described mouse line in which exon 2-6 of the *ß-catenin* gene encoding a portion of the armadillo repeat region is removed by Cre-mediated recombination, resulting in a non-functional, transcriptionally inactive protein^23,31^. In the Lmx1aCre::β-catenin LOF ChPe cell nuclei, the levels of non-phosphorylated (active) β-Catenin are nearly undetectable (Supplementary Figure S2A, B). Nuclear LEF1 protein levels decreased compared with controls (Supplementary Figure S2C, D). Furthermore, while TTR is present throughout the LOF ChPe at E16.5 and at birth (Supplementary Figure S2E, F), AQP1 is undetectable in several cells, suggesting a functional disruption of the ChPe (Supplementary Figure S2G, H).

β-CATENIN plays an important role in cell adhesion in addition to its function in transcriptional regulation. Consistent with this, both the GOF and LOF perturbations of β-CATENIN caused the ChPe to exhibit an abnormal morphology. The β-catenin GOF ChP was extensively folded, accompanied by a highly disorganized E-CADHERIN distribution (Figure S3A-B). In the β-catenin LOF ChP, even though the exons 2-6 that are deleted in the LOF allele are critical for its role in cell adhesion, the labeling pattern of E-CADHERIN appeared unaltered at E12.5. However, by birth, E-CADHERIN distribution was disorganized together with dysmorphia of the ChP, indicating that the loss of β-CATENIN may have a progressive effect on ChP morphogenesis with time in development (Figure S3C, D).

In summary, both GOF and LOF perturbations of β-CATENIN cause disruptions of the canonical Wnt pathway, with additional consequences on ChP morphology that are consistent with its role in cell adhesion. The level of nuclear β-CATENIN increased in the GOF and decreased in the LOF ChPe, and canonical Wnt targets displayed the expected effects as a result of these perturbations. We focused our subsequent analysis on the Lmx1aCre::β-catenin GOF ChPe in the mouse to examine the consequences of persistent activation of canonical Wnt signaling since this perturbation is similar to the observed pathological mutations in human β-CATENIN^20,21^. In a similar fashion, we also studied the effects of activated canonical Wnt signaling in hESC-derived organoids.

### Constitutively activated β-CATENIN causes a loss of ChPe identity

To study the effect of constitutive activation of β-CATENIN on ChP development, we first examined an age series and found a progressive loss of *Ttr* expression in the Lmx1aCre::β-catenin GOF ChPe from E12.5 to birth (Figure 3A, B). Immunostaining for apically enriched ACTIN and basal lamina marker LAMININ revealed major morphological changes from E14.5 in the form of multiple folds (Figure 3C-E). Although OTX2, a transcription factor critical for ChPe development, was detected in 93.2% of the GOF ChPe cells (compared with 98.3% in the control), quantifications indicated that its level was drastically reduced in the GOF ChPe (Figure 3F, G). AQP1 and TTR staining was undetectable in the vast majority of GOF ChPe cells, with only 2.3% or 12.4% cells positive for AQP1 or TTR, respectively, compared with 93.6% and 97.8%, respectively, in the controls (Figure 3F, H). The ventricles of Lmx1aCre::β-catenin GOF brains collapsed as development proceeded (Figure 3A), possibly due to a lack of fluid resulting from reduced AQP1 together with a reduction in several solute transporters encoded by the SLC family (*SLC12a2, 4a2, 4a10*; Figure 3I, J). The progressive nature of the phenotype is consistent with the constitutive and persistent nature of the GOF disruption of the *β-Catenin* gene.

**Figure 3:**
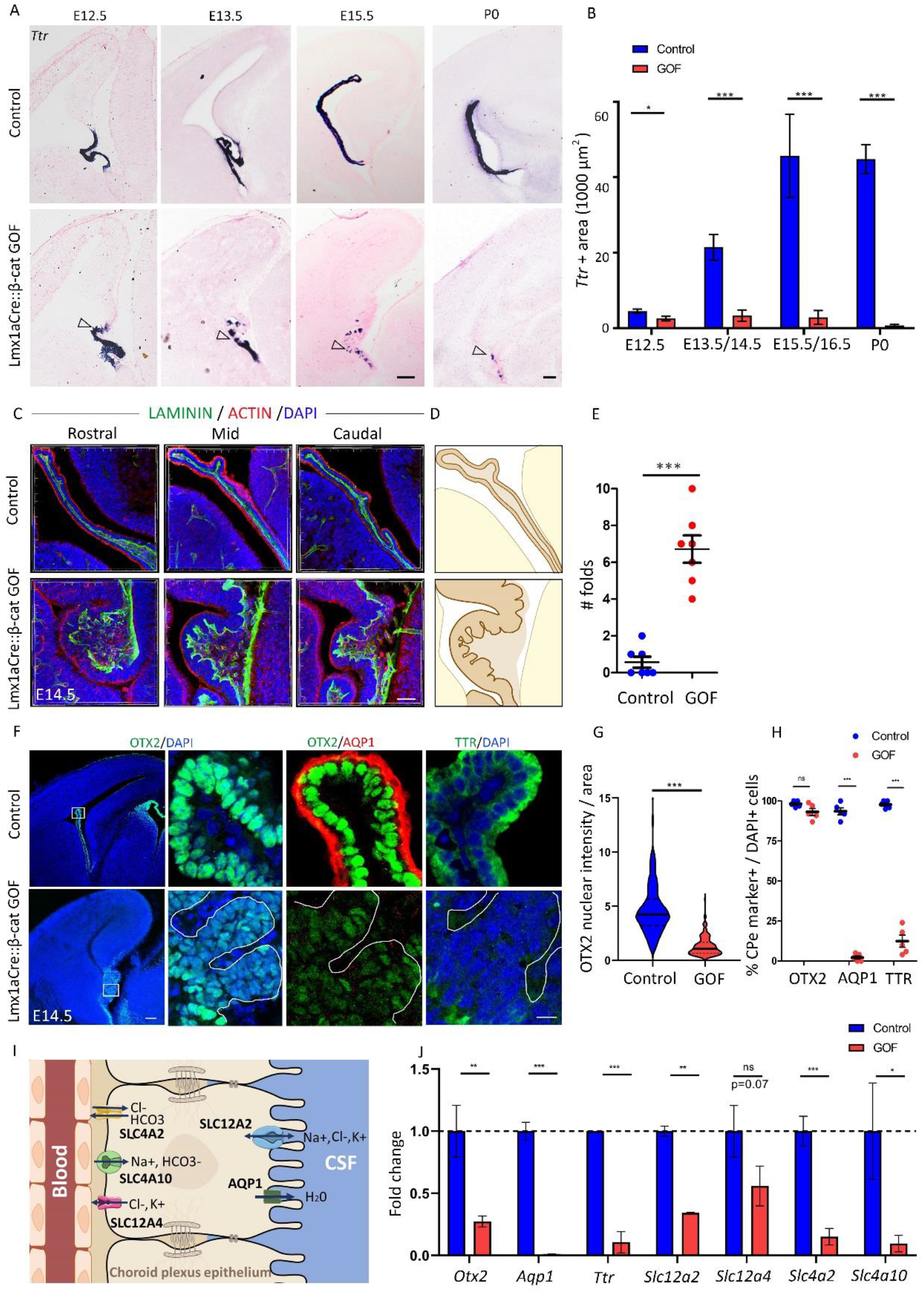
Constitutively active β-CATENIN in the hem and ChPe disrupts choroid plexus identity. (A, B) In situ hybridization for *Ttr* reveals a patchy expression that is progressively lost from E12.5-P0 in Lmx1aCre:: β-cat GOF brains compared with controls. (B) Quantification of *Ttr*-expressing area ± SEM (N=4 (E12.5), 5 (E13.5/E14.5), 4 (E15.5/E16.5) & 3 (P0). (C, D) Immunohistochemistry for ACTIN and LAMININ reveals that the ChP is extensively folded in Lmx1aCre:: β-cat GOF brains compared with controls. (E) Quantification of the number of folds (N=7). (F) ChPe markers such as AQP1 and TTR are almost undetectable, and OTX2 labeling is greatly reduced in intensity in the Lmx1aCre:: β-catenin GOF compared with controls. (G) Violin plot of OTX2 intensity/nuclear area reveals a significant reduction in the Lmx1aCre:: β-cat GOF choroid plexus (N=3). (H) Scatter plot displaying 500 DAPI+ cells scored for the co-labeling for each marker. In Lmx1aCre:: β-cat GOF ChPe 2.3% are AQP1+, 12.4% are TTR+, and 93.2% are OTX2+ whereas in control brains 93.6% are AQP1+, 97.8% are TTR+, and 98.2% are OTX2+ (N=5). (I) A cartoon illustrating transporters that are down regulated in the ChPe of β-cat GOF brains. (J) qPCR analysis of E14.5 ChP shows that not only Ttr and Otx2 but also several transporters and channels such as Aqp1, Scl12a2 (Nkcc1), Slc4a10 (NBCn2), Slc4a2 (Ae2) are significantly down regulated in the Lmx1aCre:: β-cat GOF ChP compared with control. Bar graphs represent mean ± SEM (N=3). Statistical tests in B, D, and E: two-tailed unpaired Student’s t-test with unequal variance; * p < 0.05, ** p < 0.01, *** p < 0.001, ns if p-value> 0.05. Scale bars: 100 μm (A and F low mag); 50 μm (C); 10 μm (F high mag).

### Gain of neuronal identity in the ChPe upon constitutive activation of β-CATENIN

The loss of ChPe markers in Lmx1aCre::β-catenin GOF brains prompted us to examine whether this tissue acquired a different identity. It is well established that canonical Wnt signaling from the cortical hem is necessary and sufficient for the induction of hippocampal fate in the adjacent neuroepithelium^32–34^. Therefore, we tested whether neuronal, and in particular hippocampal markers were upregulated in the Lmx1aCre::β-catenin GOF ChPe. Remarkably, at E13.5, parts of the GOF ChPe were positive for both the ChPe marker OTX2, and PAX6 which is normally restricted to neuronal ventricular zone progenitors (Figure 4A; Supplementary Figure S4). Likewise, several cells were positive for OTX2 as well as the dentate granule cell marker PROX1. Overall, 31.9% OTX2+ cells were PAX6+ and 20.2% were PROX1+ in the β-catenin GOF ChPe, compared with no co-labeling (0%) for each of these markers in controls (Figure 4A-C). The β-catenin GOF also displays enhanced proliferation as seen by PH3 labeling, and apoptosis as detected by staining for cleaved CASPASE 3 (Supplementary Figure S4).

**Figure 4.**
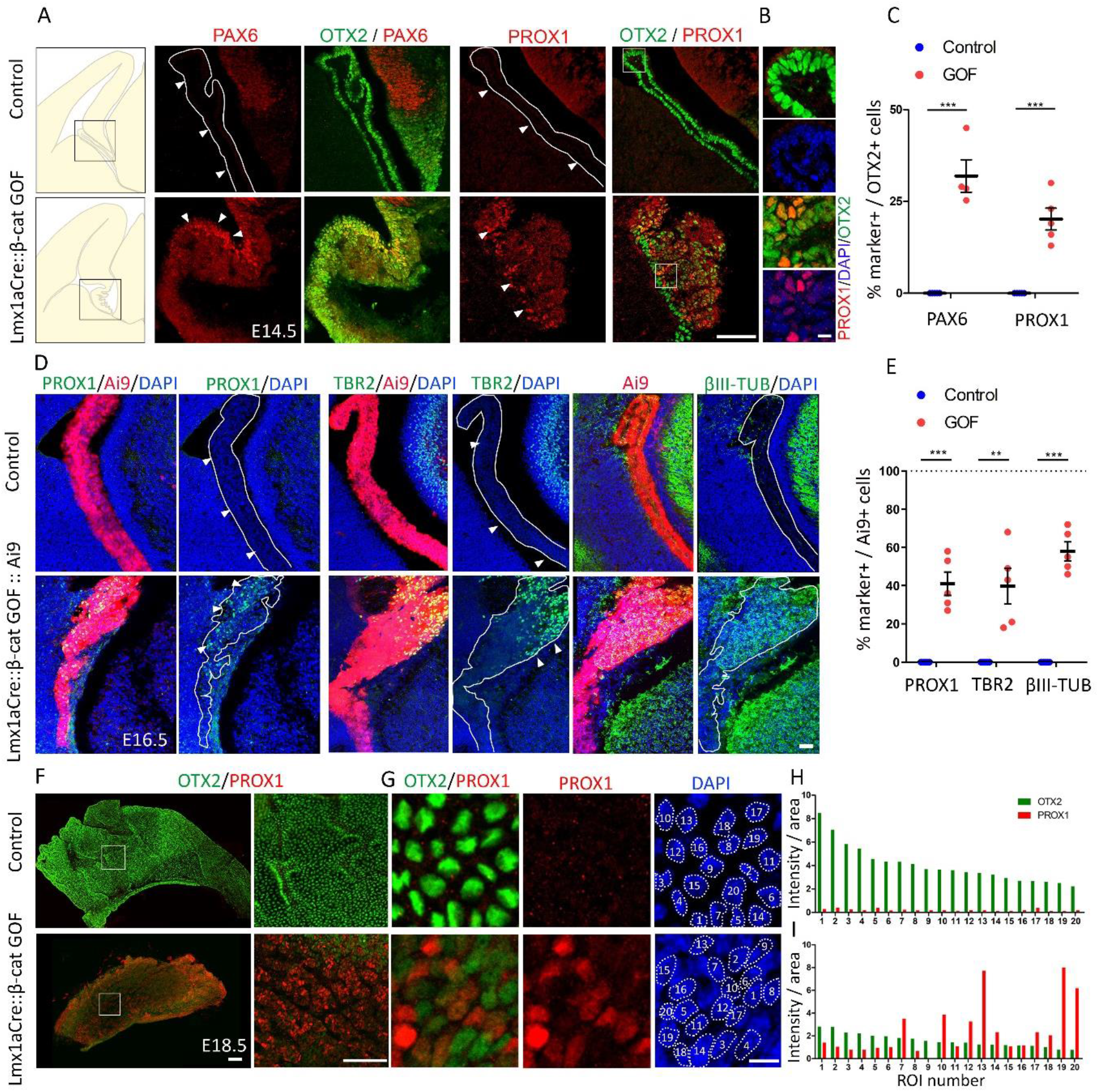
Constitutive activation of β-catenin transforms the ChPe to a neuronal and hippocampal-like molecular identity. (A-C) At E14.5, the control ChPe is positive for OTX2 but not PAX6 or the dentate granule cell marker PROX1. In the Lmx1aCre:: β-cat GOF ChPe OTX2 labeling co-localizes with that of the neuronal progenitor marker PAX6 in 31.9% (N=4) and with the granule cell marker PROX1 in 20.2% (N=5) cells. (D, E) At E16.5, using the Ai9 reporter to identify the Lmx1aCre lineage, 0% of Ai9+ control ChPe cells also label for PROX1, the intermediate progenitor marker TBR2, or the differentiated neuronal marker βIII TUBULIN. In Lmx1aCre:: β-cat GOF brains, the Ai9+ cells co-label for PROX1 (41%), TBR2 (38.9%), and βIII TUBULIN (57.9%). (F, G) Brains ChP co-immunostained for PROX1 and OTX2 using wholemount preparations from E18.5 control and Lmx1aCre:: β-cat GOF samples. The Lmx1aCre:: β-cat GOF sample displays decreased OTX2 labeling and the presence of PROX1+ cells of varying intensities which never appear in control samples. (H,I) Intensity quantification of 20 cells from a given field shows a marked decrease in OTX2 intensity in cells of the Lmx1aCre:: β-cat GOF ChPe, and high levels of PROX1 in some of these cells. Statistical tests in C and E: two-tailed unpaired multiple Student’s t-test with unequal variance; * p < 0.05, ** p < 0.01, *** p < 0.001, ns if p-value> 0.05. Scale bars: 100 μm (A and F); 10 μm (B and G); 50 μm (D).

By E16.5, 41% of Cre-expressing (Ai9) cells were positive for PROX1 (Figure 4D, E) in Lmx1aCre::β-catenin GOF brains compared with 0% in the controls. As development proceeds, the telencephalic ventricular zone progenitors give rise to a TBR2-positive pool of sub-ventricular zone progenitors. At E16.5, the GOF ChPe displayed this feature as well, such that numerous (38.9%) TBR2-positive cells were scattered in this tissue, but were never seen in controls (Figure 4D, E). In addition, the postmitotic neuronal marker βIII TUBULIN appeared in 57.9% of the Ai9+ cells in the GOF ChPe compared with 0% in controls (Figure 4D, E). The ChP is normally populated by a small number of scattered neuronal cells^35^, which also appeared in the control sections we examined. However, these do not arise from the hem and did not express Ai9, whereas the TBR2 and βIII TUBULIN positive cells in the GOF ChPe co-localized with Ai9 (Supplementary Figure S5). PROX1 labeling persisted in the Lmx1aCre::β-catenin GOF at E18.5. While PROX1 was never detected in the controls, it was present in varying intensities in the cells of the GOF ChPe in which OTX2 levels were uniformly low compared with controls (Figure 4F-I).

Lmx1aCre acts in the progenitor domain for the ChPe, the cortical hem, as well as in the differentiated ChPe^25^. We compared this line with Foxj1Cre, which acts in ciliated cells in the brain, and is known to be expressed in the primordial ChPe from E11.5^36,37^. When each Cre line was crossed to the Ai9 reporter, the differences in their expression domain were apparent: Foxj1Cre did not appear to be active at E10.5, at which stage Lmx1aCre was expressed in the hem and in the choroid plaque (arrowheads, Figure 5A, B), which is the precursor of differentiated ChPe. At E12.5, Foxj1Cre was expressed in the ChPe and by Cajal-Retzius cells, but excluded from the hem, whereas Lmx1aCre was expressed in all these structures (Figure 5A, B). Foxj1Cre::β-catenin GOF embryos displayed a dysmorphic ChP that recapitulated the loss of ChPe identity seen in Lmx1aCre::β-catenin GOF brains (Figure 5). At E16.5, only 33.2% of the Ai9 cells were also AQP1-positive and 46.8% were TTR-positive, compared with 98.5 and 98%, respectively, in the controls. While OTX2 labeling persisted in the Foxj1Cre::β-catenin GOF ChPe, its level was significantly lower than in controls (Figure 5C-E), mirroring that seen in Lmx1aCre::β-catenin GOF brains. These changes were apparent throughout the rostro-caudal extent of the Foxj1Cre::β-catenin GOF ChPe (Supplementary Figure S6). Therefore, the loss of ChPe identity in the Foxj1Cre::β-catenin GOF ChPe recapitulated that seen in the Lmx1aCre::β-catenin GOF ChPe.

**Figure 5:**
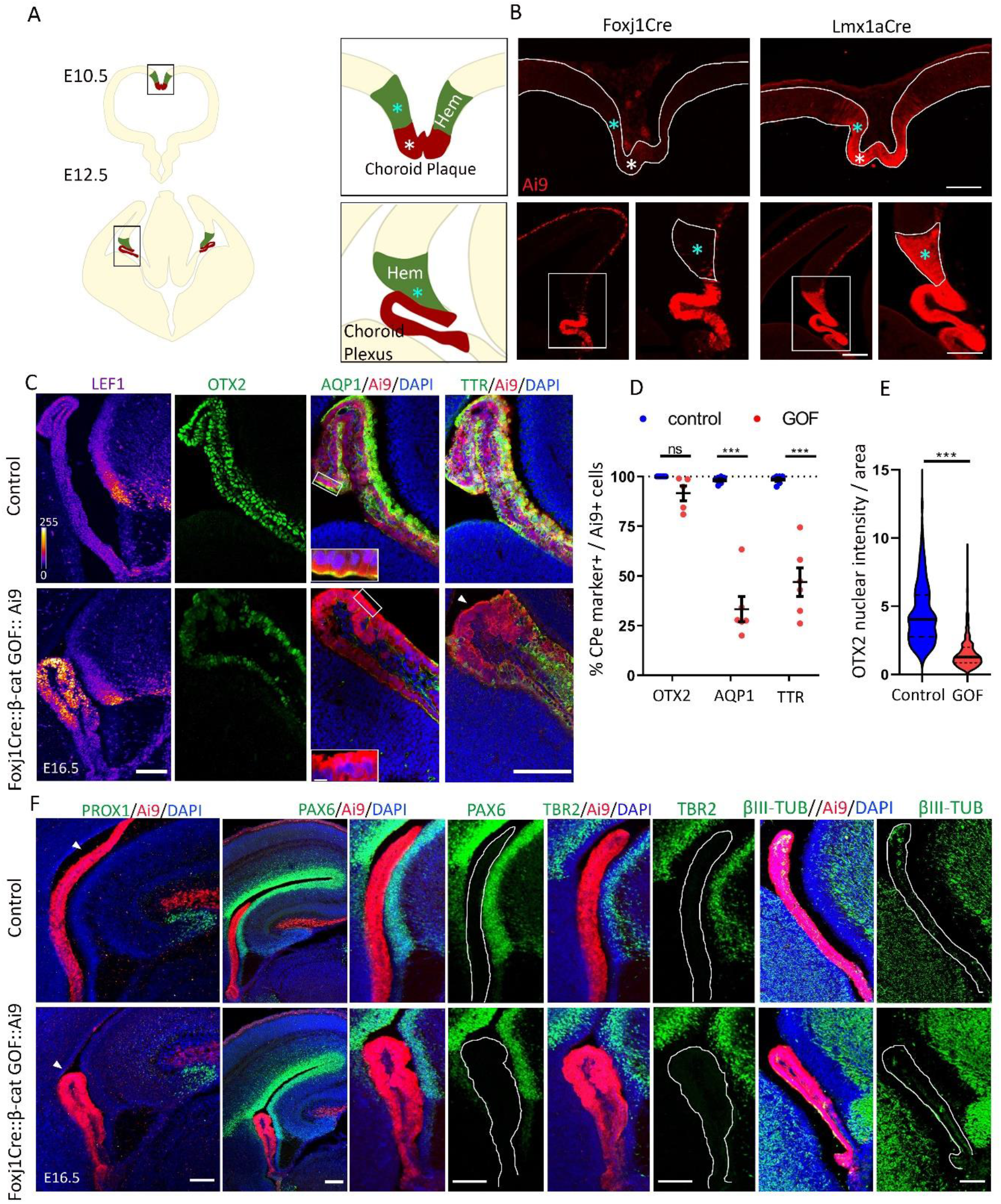
Stabilizing β-CATENIN selectively in the specified ChPe using FoxJ1Cre causes loss of ChPe fate but not transformation to a neuronal fate. (A) A cartoon illustrating the hem (cyan asterisks), choroid plaque (white asterisks), and choroid plexus at E10.5 and E12.5. (B) Foxj1Cre::Ai9 marks differentiated choroid plexus at E12.5 but not the choroid plaque at E10.5 or the hem (asterisk) at either stage. Lmx1aCre::Ai9 marks both hem and choroid plexus/plaque starting from E10.5. (C-E) In E16.5 embryos, LEF1 levels increase, while bonafide ChPe markers AQP1, TTR, and OTX2 decrease in the Foxj1Cre:: β-cat GOF compared with controls, as seen using (C) immunofluorescence and (D) quantification of the number cells positive for each ChPe marker as a fraction of the AI9-positive cells. Scatter plot showing that in the E16.5 Foxj1Cre:: β-cat GOF ChP, 33.2% of the 781 Ai9+ cells scored also display AQP1, 46.8% of 1097 cells also display TTR, and 91.7% of 731 also display OTX2 labeling (N=5 for OTX2 and N=6 for TTR & AQP1). (D, E) However, the nuclear intensity of OTX2 decreases significantly in Foxj1Cre:: β-cat GOF ChPe cells compared with controls (N=5; 536 nuclei from control and 590 nuclei from Foxj1Cre:: β-cat GOF ChPe). Statistical test: two-tailed unpaired multiple Student’s t-test with unequal variance; * p < 0.05, ** p < 0.01, *** p < 0.001, ns if p-value> 0.05. (F) Presence of the granule cell marker PROX1 is seen in the dentate gyrus, neuronal progenitor markers PAX6 and TBR2 appear in the telencephalic ventricular and sub-ventricular zones, respectively, and βIII-TUBULIN is present in postmitotic neurons in the control and Foxj1Cre:: β-cat GOF dorsal telencephalon. None of these markers appear in the ChPe of either genotype, except for a small number of neurons that normally populate the ChPe and are βIII-TUBULIN positive. Scale bars: 100 μm (B, C & F); 10 μm (C inset).

In contrast, the Foxj1Cre:: β-catenin GOF ChPe failed to display any transformation to a neuronal fate. None of the markers we reported in the Lmx1aCre::β-catenin GOF ChPe, i.e. PROX1, PAX6, TBR2, or βIII TUBULIN, were detected in the Foxj1Cre:: β-catenin GOF ChPe (Figure 5F). This offers an insight into the timing and stage at which the canonical Wnt pathway can modulate neuronal fate specification in cell types that normally take on ChPe fate.

Since loss of ChPe identity was accompanied by a gain of neuronal identity only when the β-catenin GOF was initiated in the hem, it was important to examine whether ChPe progenitors were converted to neuronal progenitors or whether specified ChPe cells that displayed ChPe markers were transformed to a neuronal fate. Therefore, we examined whether neuronal markers appear in cells that also display ChPe markers. At E12.5, both control and Lmx1aCre::β-catenin GOF brains displayed *Wnt2b* and BLBP restricted to the hem while *Ttr* mRNA and TTR protein was restricted to the ChPe (Figure 6A, B). Whereas βIII TUBULIN staining was almost undetectable in control ChPe, a small region of the β-catenin GOF ChPe was βIII TUBULIN positive, co-localizing with TTR immunostaining (Figure 6C). This indicated that neuronal markers were upregulated in specified ChPe cells from early stages in its development.

**Figure 6:**
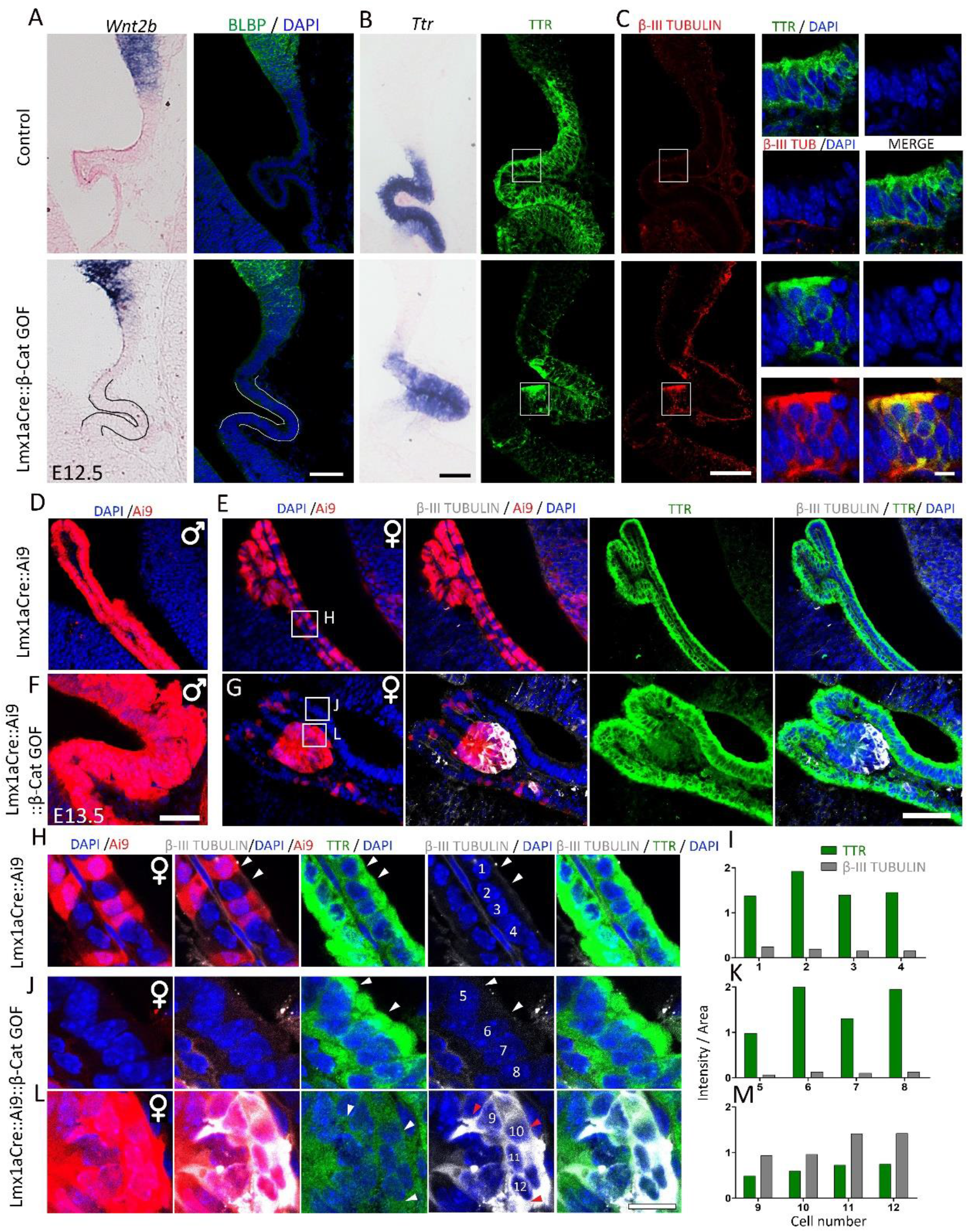
β-Catenin GOF causes upregulation of β-III TUBULIN in TTR-positive ChPe cells. (A-C) At E12.5, both control and Lmx1aCre::β-Catenin GOF ChP display a ribbon-like morphology and do not express hem *Wnt2b* and BLBP(A) but express *Ttr* mRNA and TTR protein (B). (C) Co-immunostaining for β-III TUBULIN and TTR identifies cells that display both markers selectively in the β-Catenin GOF ChPe. (D-G) At E13.5, male Lmx1aCre::Ai9 control and Lmx1aCre::Ai9::β-Catenin GOF brains display Cre-driven Ai9-positive cells throughout the ChPe (D, F). In contrast, female control and β-Catenin GOF brains show mosaic Ai9 due to random X-inactivation. β-III TUBULIN is undetectable in the female control ChPe, but appears selectively in Ai9+ patches in the female β-Catenin GOF ChPe (E,G). (H-K) β-III TUBULIN is undetectable and therefore does not co-localize with TTR staining in Lmx1aCre::Ai9 female control ChPe cells (cells #1-4; H,I) and in non-Ai9 cells internal control cells in Lmx1aCre::Ai9::β-Catenin GOF female ChPe (cells #5-8; J,K). These markers are both detected in Ai9-positive Lmx1aCre::Ai9::β-Catenin GOF female ChPe cells (#9-12; L, M). (I, K, M) Fluorescence intensity quantifications of cells #1-12 from (H,J,L). Scale bars: 10 μm (C high mag & H-L); 50 μm (A-E).

To examine this neuronal transformation further, we took advantage of X-inactivation-based mosaic expression of the Cre recombinase in Lmx1aCre female embryos. Whereas male brains expressed the Ai9 reporter in the entire *Lmx1a* expression domain (Figure 6D, F), female Lmx1aCre brains displayed a mosaic expression of the Ai9 reporter (Figure 6E, G), in both control (Lmx1aCre::Ai9) and Lmx1aCre::Ai9::β-catenin GOF animals. In control female brains, all ChPe cells, whether Ai9 positive and negative, were TTR positive but did not display detectable βIII TUBULIN immunofluorescence (Figure 6E, H). In β-catenin GOF female brains, ChPe cells that were Ai9-negative (no Cre activity) were indistinguishable from controls (Figure 6G, J). Consistent with the findings in male β-catenin GOF brains, some Ai9-positive (Cre-active) cells in the ChPe of female β-catenin GOF brains displayed βIII TUBULIN immunofluorescence in cells that were also TTR positive (Figure 6G, L). Similar results were seen when ChPe marker E-CADHERIN was examined together with βIII TUBULIN in female β-catenin GOF brains (Supplementary Figure S7A): ChPe cells that were Ai9-negative (no Cre activity) never displayed βIII TUBULIN staining (Supplementary Figure S7B); whereas some Ai9-positive (Cre-active) cells were positive for both E-CADHERIN and βIII TUBULIN labeling (Supplementary Figure S7C). In summary, the mosaic expression of the Cre recombinase in female embryos offered an elegant internal control in which Cre-driven β-catenin GOF cells resided adjacent to control (no Cre activity) cells in the same ChPe tissue. These data demonstrated that ChPe cells that were TTR or E-CADHERIN positive also expressed βIII TUBULIN in β-catenin GOF brains, and therefore led to the conclusion that cells that were specified as ChPe became transformed to a neuronal fate. The observation that βIII TUBULIN positive cells were always Ai9 positive further suggests the effect of β-catenin GOF was likely to be cell-autonomous.

To examine the nature of this apparent neuronal transformation of the Lmx1aCre::β-catenin GOF ChP in a comprehensive manner, we compared transcriptomic datasets obtained from micro dissected E14.5 hippocampal primordia isolated from the same telencephalic hemispheres from which the ChP was isolated for RNAseq (Figure 2), as illustrated in the cartoon (Figure 7A; Supplementary Data file). Principal Component Analysis (PCA) of the overall transcriptomes revealed that the Lmx1aCre::β-catenin GOF ChP is more similar to the control and GOF hippocampus than it is to the control ChP (Figure 7B). Consistent with this, the expression of genes critical for ChP development and function such as *Aqp1, Htr2c, Kcne2, Otx2, Slc12a2(Nkcc1),* are dramatically reduced, and the expression of genes important for neuronal development such as *Emx1, Emx2, Fezf2, Neurod1, Prox1* are up-regulated in the β-catenin GOF ChP (Figure 7C). Hierarchical clustering of the top 500 genes expressed in the ChP and the hippocampus recapitulates this loss of choroid epithelial fate and gain of neuronal fate in the Lmx1aCre::β-catenin GOF ChP (Figure 7D; Supplementary Data file).

**Figure 7.**
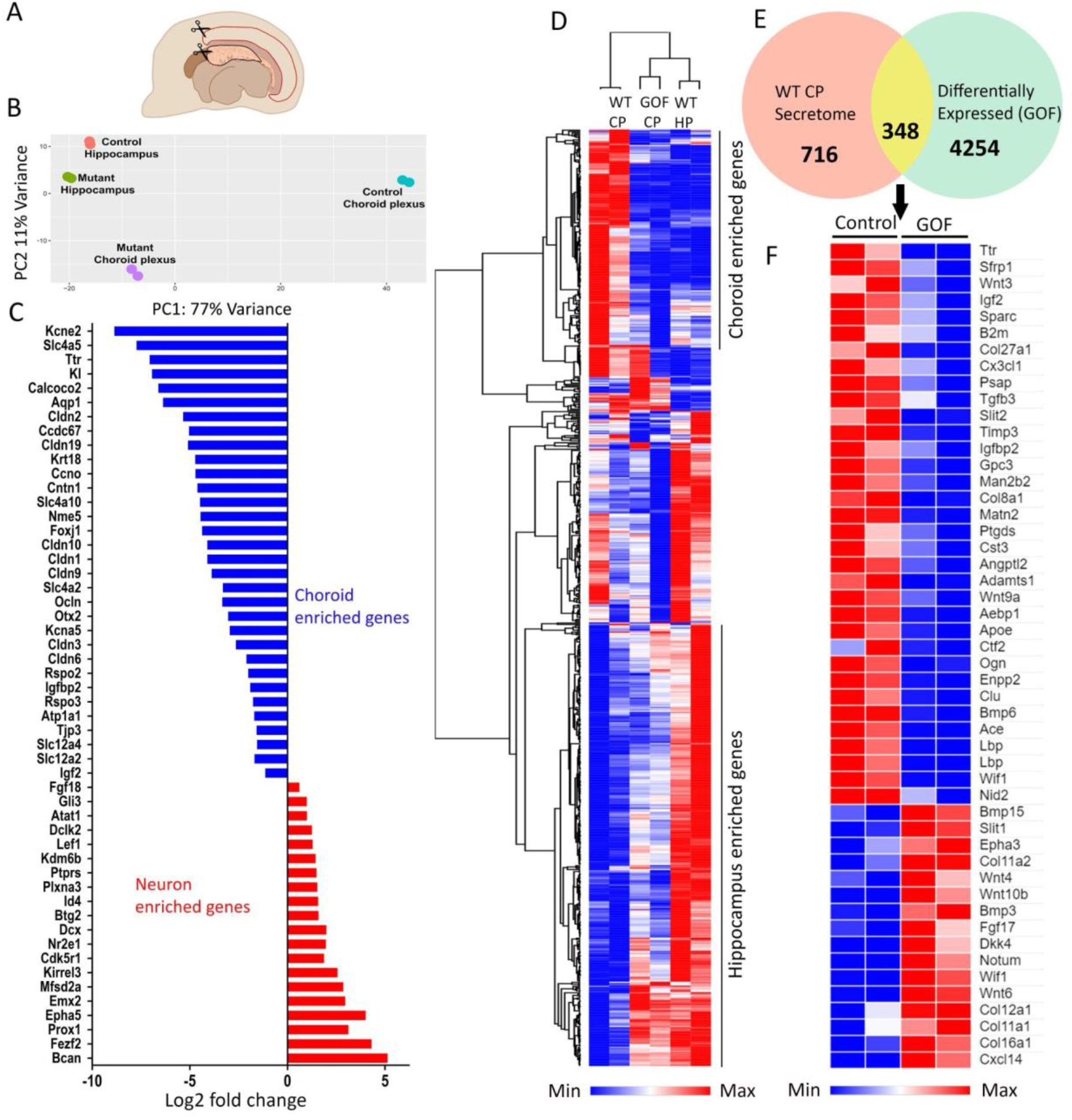
Constitutive activation of β-CATENIN catenin leads to a dysregulation of the ChP transcriptome. (A) A cartoon depicting the microdissection E14.5 ChP and hippocampus for RNAseq analysis. (B) Principal Component Analysis of the RNAseq datasets. (C, D) Choroid plexus-enriched genes are downregulated, and hippocampus-enriched genes are upregulated in the Lmx1aCre:: β-catenin GOF ChP (E, F) Venn diagram showing the overlap between genes corresponding to the ChP secretome (from Lun et al. 2015.) and differentially expressed genes in the Lmx1aCre:: β-catenin GOF versus control ChP and heat map of selected genes from (E) shows the dysregulated genes belong to various known pathways of cell-cell signalling (F).

A recent study used single cell RNAseq for transcriptomic analysis of the developing mouse ChP and identified five distinct cell types in this tissue, namely epithelial, neuronal, endothelial, mesenchymal, and immune cells. Dani et al., 2021^35^ identified 86 ChP epithelial cell-enriched genes in the ChP, and of these, 74 (86 %) were significantly downregulated in the Lmx1aCre::β-catenin GOF ChP, consistent with our interpretation of a loss of ChPe fate; none were upregulated; 1 gene did not change in expression, and 11 were not detected. In contrast, of the 274 neuronal-enriched genes, 123 (45 %) were upregulated in the Lmx1aCre::β-catenin GOF ChP, consistent with our interpretation of a gain of neuronal fate; 21 were downregulated and 122 did not change in expression; 9 were not detected (Figure 8 A-C; Supplementary Data file). The genes upregulated in the GOF ChP included several known regulators of neuron differentiation such as bHLH members *Neurod1, Neurod2, Neurod6 Neurog2*, *Tcf3*; *Sox21, Sox4, Sox1, Zeb1* and *Zeb2* (Figure 8C, Supplementary Figure S7).

**Figure 8:**
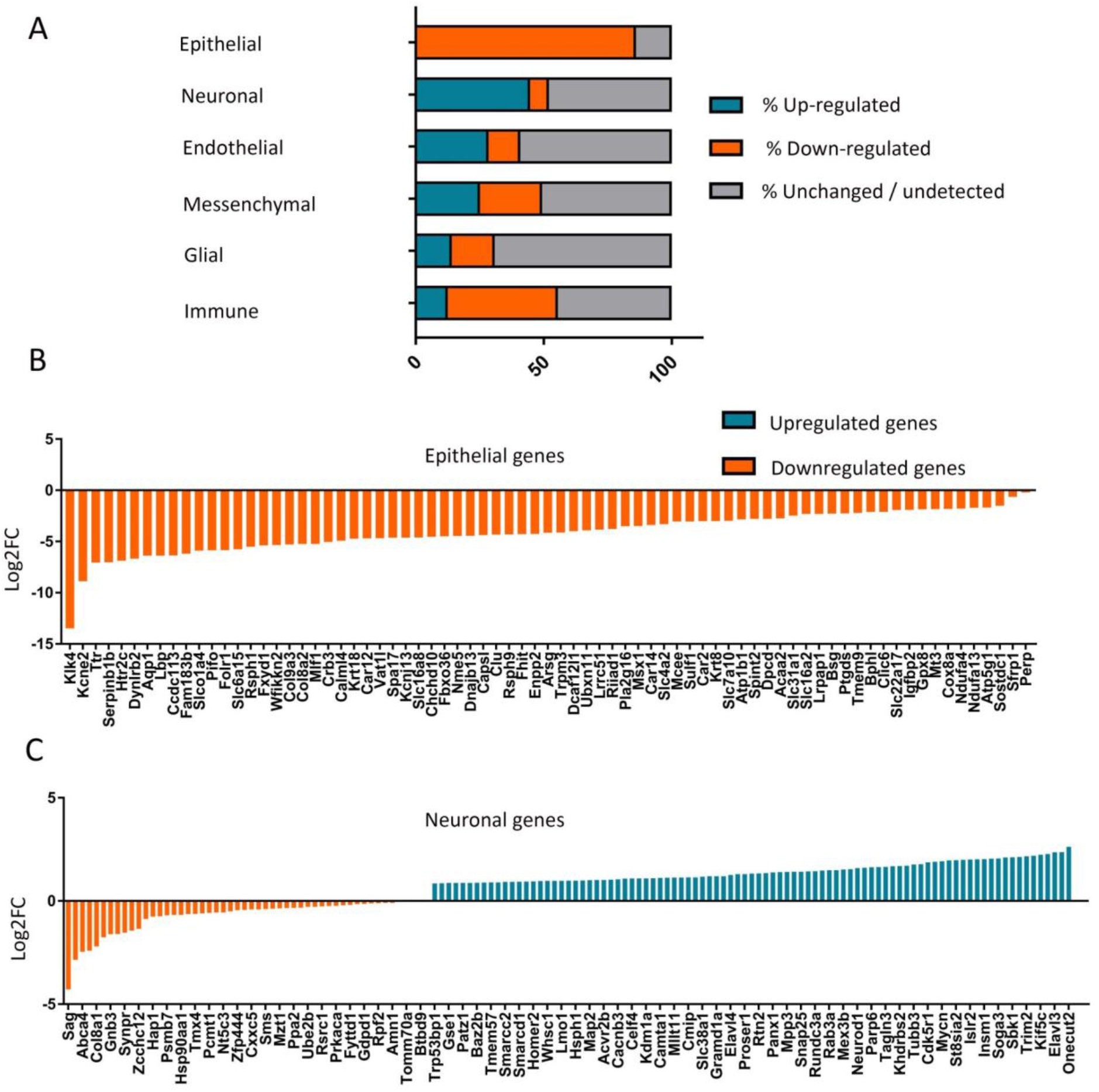
(A) comparison of RNA-seq data from E14.5 Lmx1aCre:: β-catenin GOF ChP with scRNAseq data from E16.5 ChP (Dani et al., 2021). Genes that were differentially expressed (DE) between the Lmx1aCre:: β-catenin GOF ChP and control ChP (Supplementary Data file) were compared with the genes enriched in epithelial/neuronal/endothelilal/mesenchymal/glial/immune cell groups identified in Dani et al. (2021). For each cell group, the DE list contained genes that were either upregulated (green), downregulated (orange), or unchanged/undetected (grey). (B, C) Selected epithelial- and neuronal-enriched genes (Dani et al., 2021) that were dysregulated in the Lmx1aCre:: β-catenin GOF ChP.

Although the remaining cell types identified in Dani et al., 2021^35^ are not known to arise from the hem, it is intriguing that a substantial fraction of genes enriched in these populations were also dysregulated in the Lmx1aCre::β-catenin GOF ChP, suggesting the possibility of non-cell autonomous effects of β-catenin GOF in the hem-derived cells. This further suggests interactions between these cell types that have thus far not been examined.

Consistent with the apparent neuronal transformation of the GOF ChP, gene ontology analysis of the Lmx1aCre::β-catenin GOF transcriptome indicated that genes that were upregulated in the GOF ChP belonged to the categories like neurogenesis (GO: 0022008), neuron projection development (GO: 0031175), cell differentiation (GO: 0030154), dentate gyrus development (GO: 0021766, GO: 0021542). We also examined GO categories in which genes were downregulated in the GOF ChP, and found that these belonged to classes important for ChP function such as cilium organization (GO:0044782) and ion/solute transport (Supplementary Figure S8D). We examined the effects of this transformation on the CSF in Lmx1aCre::β-catenin GOF brains. It was not possible to extract CSF from Lmx1aCre::β-catenin GOF brains because the ventricles were collapsed from E13.5 (Figure 3A). Therefore, we took the alternative approach of examining a curated list of 1064 ChP-secretome genes from the published literature^38^ and comparing it with the set of genes differentially expressed in the GOF versus control ChP. 348 of the ChP secretome genes were dysregulated in the GOF ChP (Figure 7E). The majority (211/348) of these genes were downregulated compared with the control ChP, including several major components of the CSF such as *Igf2, Igfbp2, Ttr* (Figure 7F). Some genes were upregulated in the GOF ChP, such as *Bmp6, Fgf17, Wnt4&6,* suggesting a change in the composition of the secreted CSF in Lmx1aCre::β-catenin GOF brains. Together these results suggest that the ChP has partially transformed, acquired neuronal characteristics, and the gene expression data suggest an altered secretome that would impact CSF composition.

### ChPe-like structures arise in hESC organoids upon a combination of BMP and canonical Wnt pathway activation

We sought to examine whether constitutive activation of canonical Wnt signaling disrupts ChPe identity in the human, as it does in the mouse. First, we used established protocols to achieve ChPe differentiation in hESC-derived organoids^39^. In this protocol, hESCs are aggregated and seeded in a 96 well low-adhesion plate, and over 18 days of incubation, they form spherical structures which grow in diameter (Figure 9A, B). At this stage, a morphogen cocktail containing BMP4 and CHIR, a GSK3 inhibitor, is added. GSK3 is responsible for phosphorylating β-CATENIN and marking it for degradation, so CHIR addition results in a greater amount of active β-CATENIN^40^. However, unlike the β-catenin GOF mouse, CHIR can be added in graded amounts to control the extent of canonical Wnt pathway activation^41^. A cocktail containing a low level of CHIR (3 μM) together with BMP4 (0.5nM) promotes ChPe-like differentiation^39^. The spherical organoids became slightly tapering and developed multiple protrusions within a few days of exposure to the cocktail (Figure 9B). Immunohistochemistry revealed ChPe markers TTR, AQP1, and OTX2 were present along the perimeter of the protrusions, with cells displaying a cuboidal morphology (Figure 9C-E). To better characterize these organoids, we performed RNA-seq after 12 days of exposure to the cocktail. Even though the ChPe-like differentiation was seen only along the perimeter of the organoid, the transcriptomic analysis of the sample revealed characteristic markers of the ChPe such as *AQP1, CLIC6, KRT18 and TTR* (Figure 9F). Interestingly, genes associated with the blood-CSF barrier (BCSFB) such as *CDH1*, *CLDN3*, *JAM2, TJP1, and TJP2* and were also upregulated, as were several transporters and ion channels encoded by the SLC gene family and genes associated with ciliogenesis, consistent with the ciliated epithelium nature of the ChPe (Supplementary Figure S9). In summary, consistent with Sakaguchi et al. (2015)^39^, ChPe-like structures develop in organoids upon exposure to a cocktail of BMP4 and low levels of CHIR.

**Figure 9:**
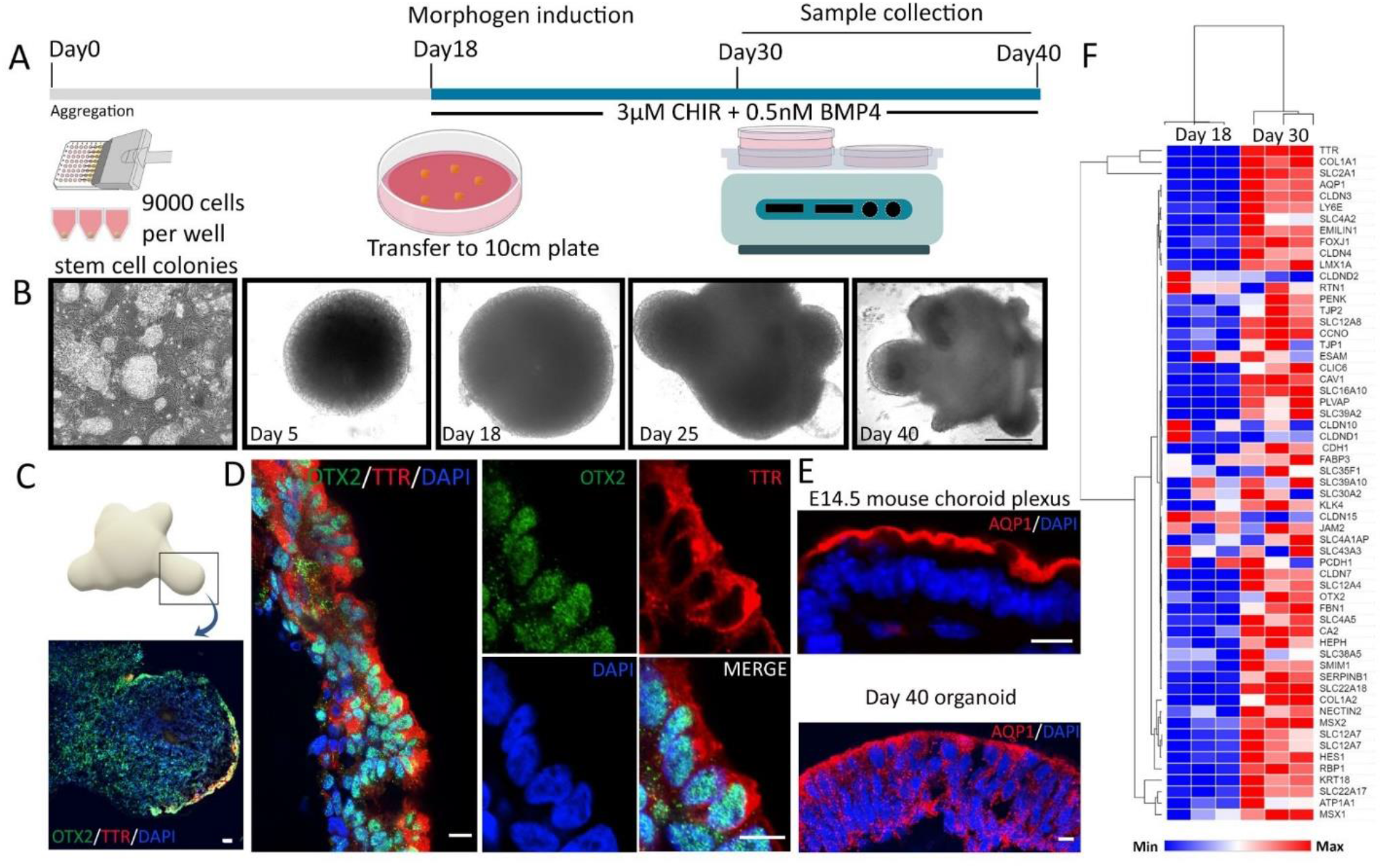
Differentiation of hESC organoid cultures using a ChPe differentiation-promoting protocol. (A) Schematic presentation describing the organoid culture protocol. (B) Bright-field images of organoids at different stages of development. (C,D) Immunostaining reveals the presence of choroid markers OTX2 and TTR in the perimeter of the protrusions of a day 40 organoid. (E) AQP1 immunostaining in Day 40 organoids is similar to that seen in the E14.5 mouse ChP. (F) Heat map generated from normalized reads obtained by MARS-Seq shows upregulation of ChP-enriched genes by day 30 of the organoid culture. Scale bars: 200 μm (B); 100 μm (C); 10 μm (D and E).

### Loss of ChPe-like identity upon increased activation of canonical Wnt signaling in hESC-derived organoids

We examined the effect of increased activation of canonical Wnt signaling in the hESC-derived organoids. We tested a range of concentrations of CHIR (3 μM to 12 μM) in the cocktail added to the organoids after 18 days *in vitro*, keeping the BMP4 concentration constant. The organoids were examined for the presence of *Ttr/* TTR, *Aqp1*/AQP1, and *Otx2*/OTX2 by immunohistochemistry, qPCR analysis, and RNA-Seq (Figure 10). As early as 7 days post CHIR treatment (25 days *in vitro*), 3 μM-treated organoids displayed multiple domains of TTR decorating the perimeter, but 12 μM-treated organoids displayed no detectable TTR (Figure 10C; Supplementary Figure 10C). We tested the dose-response to CHIR treatment by immunostaining for TTR, AQP1, and OTX2 in organoids using the different CHIR regimens. Whereas all three proteins appeared along the perimeter of 3 μM CHIR-treated organoids, they were undetectable in organoids treated with higher CHIR concentrations (Figure 10D; Supplementary Figure S10D). qPCR analysis confirmed this finding: upon treatment with 3 μM CHIR, *TTR, AQP1,* and *OTX2* displayed a significant increase in fold-expression over the baseline (untreated 18-day organoids). However, treatment with 12 μM CHIR appeared to suppress the expression of these genes to levels below the baseline (Figure 10E). In contrast, *LEF1* expression correctly reflected the increased activation of canonical Wnt signaling by displaying a graded increase in fold expression above baseline upon 3 μM and 12 μM CHIR treatment (Figure 10F).

**Figure 10.**
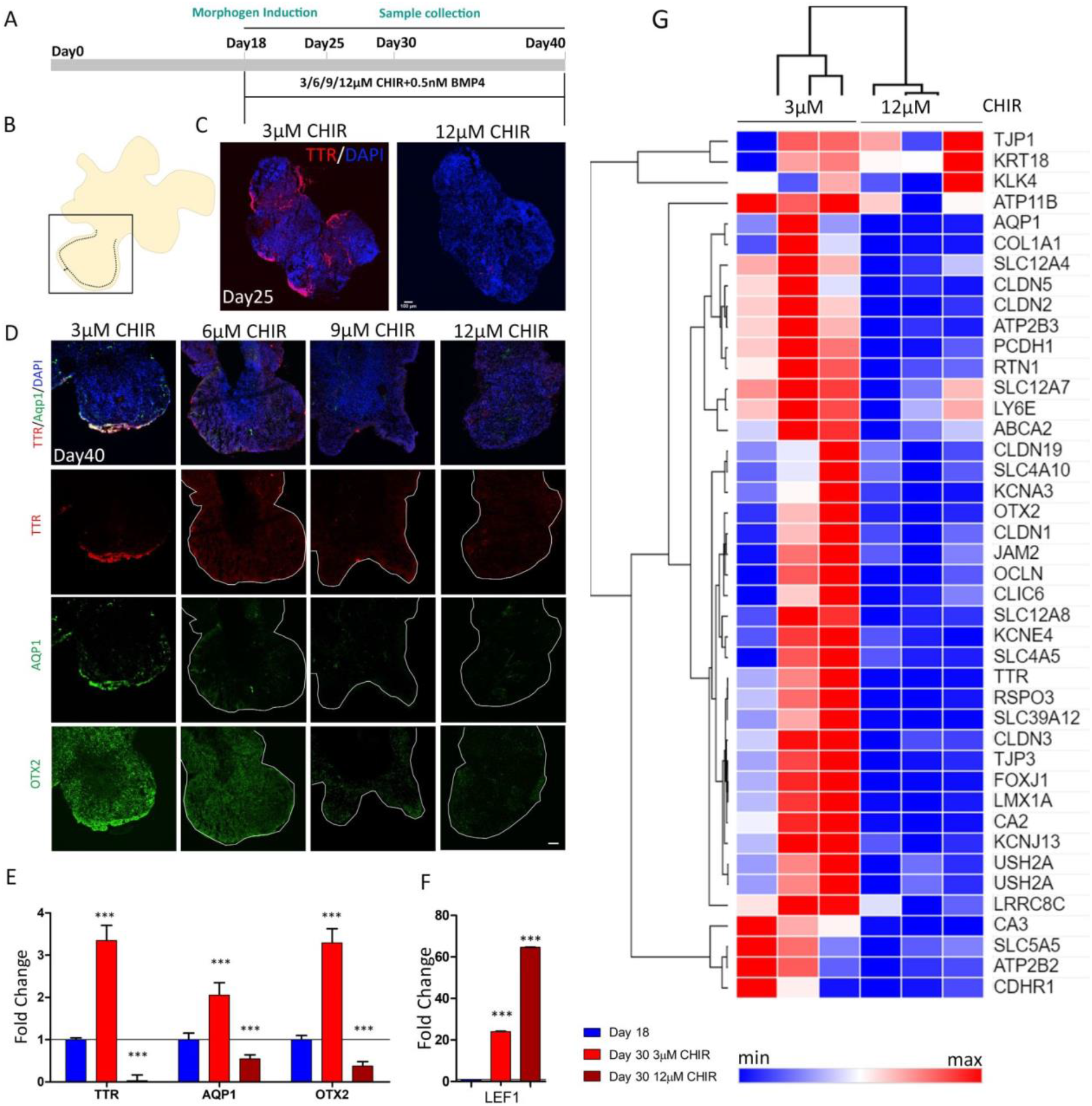
High activation of the canonical Wnt pathway in hESC-derived organoids causes suppression of ChPe markers. (A) A schematic presentation of the protocol for the treatment of organoids with the canonical Wnt agonist CHIR. (B, C) The ChP marker TTR is present in the periphery of a day 25 organoid exposed to 3 μM, but not when exposed to 12 μM CHIR, from day 18. (D) High magnification images (boxed region shown in B) of serial sections of a day 40 organoid exposed to 3 μM reveal the presence of TTR, AQP1, and OTX2 in the periphery. This labeling is reduced or undetectable upon exposure to higher concentrations of CHIR (6/9/12 μM). Additional examples are in Supplementary Figure S10. (E) q-PCR analysis shows that the expression of the ChP markers *TTR, AQP1,* and *OTX2* is upregulated in organoids at day 30 organoids exposed to 3 μM CHIR from day 18. This increased expression is lost upon treatment with 12 μM CHIR for the same period. (F) *LEF1* expression displays a stepwise increase upon 3 μM and 18 μM CHIR treatment, consistent with increased activation of the canonical Wnt pathway. Group means were compared using one-way ANOVA followed by posthoc Tukey’s multiple comparison test, * p-value<0.05, **p-value<0.01, *** p-value<0.001 (N=4). (G) Heatmap comparing normalized RNA-seq reads for 3 μM and 12 μM CHIR-treated organoids. Several ChP-enriched genes are downregulated with increased activation (12 μM CHIR) of canonical Wnt signaling. All scale bars: 100 μm.

To obtain a comprehensive picture of the effect of increased canonical Wnt pathway activation on hESC-derived organoids, we performed RNA-seq of 3 μM and 12 μM CHIR-treated samples. Several major ChPe markers detected in the 3 μM-treated condition are suppressed upon 12 μM CHIR treatment (Figure 10G). These include markers of differentiated ChPe such as *AQP1, TTR, OTX2*; genes that encode proteins that participate in forming tight junctions that are typical of the ChPe, such as *CLDN1, 2, 5,* and *TJP1,3*; and those that encode transporters or ion channels such as *CLIC6, SLC4A10, SLC12A7, and SLC12A8.* The transcriptome of the 12 μM CHIR-treated organoids did not display upregulation of neuronal markers. One possible reason for this could be that any such transformation would have been limited to the periphery of the organoid, and the corresponding transcriptomic changes may have become masked due to the larger contribution of the rest of the organoid. We, therefore, examined 3 μM, 6 μM, 9 μM, and 12 μM CHIR-treated organoid samples by immunohistochemistry using a neuronal marker that is enriched in the hippocampus, ZBTB20^42^ and a choroid epithelium marker OTX2. We discovered that with increasing CHIR treatment, the organoid samples reveal a significant reduction in OTX2+ cells and a corresponding increase in ZBTB20+ cells (Supplementary Figure S11). Additional analysis would be needed to test whether this indeed represents a transformation to a neuronal identity. In summary, stabilizing β-CATENIN in hESC-derived organoids leads to the loss of ChPe identity, consistent with the findings in the β-Catenin GOF mouse.

## Discussion

The ChP actively processes canonical Wnt signaling from the earliest stages of its development. Expression of the reporter BAT-Gal and target gene *Axin2* was reported in the choroid plaque at E10.5 and the ChPe at E11.5^34^. Similarly, the expression of both *Lef1,* as well as a LEF1-βGal allele, were reported in the E12.5 mouse ChP^43^. Furthermore, the expression of *Lef1, Axin2*^43,44^, *Tcf7* (named *Tcf1* in^45^), several Frizzled receptors (*Fzd1, Fzd2, Fzd3*, and *Fzd7*)^46^ as well as the phosphorylated LRP6 co-receptor is present in the developing ChPe^47^. Consistent with these reports, we found that loss of β-CATENIN, which mediates canonical Wnt signaling, disrupts the normal development of the telencephalic ChP.

It appears, however, that a regulated level of canonical Wnt signaling is critical for ChPe differentiation. Constitutive activation of canonical Wnt signaling results in a profoundly defective ChPe both in the mouse embryo and in hESC-derived ChP organoids. Since this pathway regulates multiple aspects of cellular, tissue, and organ development, several other systems that require canonical Wnt signaling for normal development also display an array of defects upon its constitutive activation. During human hair follicle morphogenesis, loss or constitutive activation of β-CATENIN leads to tumor formation or defective differentiation, respectively^48,49^. There are some examples in which constitutive activation of β-CATENIN appears to transform cell identity. The mouse epiblast prematurely displays epithelial to mesenchymal transition^50^; endothelial cells of the midbrain transform to a “barrier cell” identity^51^, and airway epithelial cells take on a neuroendocrine-like fate^52^. Our study reports an apparent transformation of specified ChPe cells to a neuronal fate upon constitutive activation of β-CATENIN in their progenitor domain, the hem, which parallels the findings in the other systems mentioned above.

An unexpected feature is that the transcriptome of the β-catenin GOF ChP has some components that fit with a CR cell-like profile, such as *Ebf2*, *Jph3, Reelin*, *Slit1*^53^ and others that fit with a medial cortical/hippocampal-like profile *(Lef1, Prox1, Wif1)*, in addition to an array of cortical neuroepithelial or postmitotic neuronal markers (Supplementary Table 1). While it is not possible to delineate whether there is a heterogenous population of CR-like, hippocampal-like, and cortical neuron/neuronal progenitor-like cells in the β-catenin GOF ChP using bulk RNAseq, it appears that the overall picture is one of mixed transcriptomic signatures, at least at a population level. Fate choice between ChPe and CR cells is regulated by transcription repressors of the bHLH family members, HES1, 2, and 3, that promote epithelial identity by suppressing neurogenic genes and inducing BMP signaling^54^. Upon loss of all three *Hes* genes, the hem upregulates the pro-neuronal factor *Neurog2* and produces CR cells instead of ChPe cells^55^. Our results suggest that the canonical Wnt pathway may intersect with this regulatory mechanism. β-CATENIN is also involved in cell adhesion, which is an important factor in epithelial organization. In addition, the hem also expresses *Wnt5a*^12^ and WNT5a acts via the non-canonical Wnt-PCP pathway to regulate ChPe cytoarchitecture, apicobasal polarity, and overall size and complexity of the ChP^18^. BMPs are known to be critical for the specification of the ChP^16^. Thus, multiple signaling molecules of the WNT and BMP families present in the hem are each critical for ChP development. A comprehensive analysis of interactions between these factors and the pathways they regulate may offer insight on the regulation of multipotency of progenitors in the hem, and the temporal trajectory of ChPe differentiation.

Activation of the canonical Wnt pathway is often tumorigenic^21^. Human cases of ChP carcinoma are associated with mutations in exon 3 of the *ß-CATENIN* gene that lead to its constitutive activation^20,21^. Mouse models of ChP papilloma that are generated by constitutive activation of NOTCH signaling show concomitant upregulation of the canonical Wnt pathway^22^. It is not clear what mechanisms control whether canonical Wnt activation causes cell fate transformations or tumorigenic changes, but it is likely that crosstalk with other major signaling pathways is involved. The β-catenin GOF mouse ChP offers a useful and accessible model in which this question can be examined.

Key features of β-catenin GOF in the ChP include dysregulation of factors that are essential for the ChP to produce the CSF. These include several anion cation exchangers, ion channels, and solute transporters, each of which acts as a regulator of CSF composition: a sodium-potassium ATPase, *Atp1a1*^55^; an epithelial sodium channel, *Scnn1a*^56^; an inward rectifying potassium channel, *Kcnj13*^57^; a voltage-gated potassium channel, *Kcna1*^58^; and a sodium-potassium-chloride cotransporter 1, *Nkcc1*, that modulates water transport underlying CSF production^59^ and is also associated with Blood CSF barrier disruption^60^. Furthermore, the expression of several genes encoding for glucose, chloride, and amino acid transporters, is also downregulated in the β-catenin GOF ChP. Together, the data suggest that canonical Wnt signaling may be an important factor in enabling the ChPe to produce normal CSF. Mutations in exon 3 of *ß-CATENIN* in humans result in stabilization of β-CATENIN^61^, similar to that seen in the β-catenin GOF mouse model^26^. Human studies have focused on the association of these mutations with carcinomas^21,61^, but no associated effects on the CSF have been explored. Our study identifies a potentially important developmental outcome of the β-catenin GOF perturbation in terms of its possible effect on the composition of the CSF. The formation of ChPe in hESC-derived organoids requires low activation of the canonical Wnt pathway together with BMP signaling. Loss of ChPe fate in these organoids upon high activation of the canonical Wnt pathway indicates conservation of the role of this pathway in the development of the ChPe. hESC-derived organoids are not only an excellent system to study conserved developmental mechanisms but also offer the opportunity to test for subtle effects of altered ChP/CSF status on neuronal cells. Depending on the protocol used, the organoids develop into fluid-filled cysts that contain several human CSF proteins^62^. These organoids are permeable to small molecules and are excellent model systems to test modalities of drug delivery to the CNS^62^. Thus, hESC-derived ChP organoids offer a suitable system to model both diseases as well as modalities for treatment.

In conclusion, we demonstrate that the developing choroid plexus expresses components of the canonical Wnt signaling pathway in embryonic mouse and human brains. Both loss and gain of function of β-CATENIN disrupt ChPe development, indicating that this pathway is likely to be under a finely tuned regulation *in vivo*. These findings motivate future studies on how this regulation is achieved and may provide a mechanistic basis for subtle deficits in ChP/CSF function connected to the disruptions of the canonical Wnt pathway function in this system.

## Material and methods

### Tissue collection

#### Mice

All animal protocols were approved by the Institutional Animal Ethics Committee of the Tata Institute of Fundamental Research, Mumbai, India. The mouse lines used in this study were kind gifts from Kathy Millen (Center for Integrative Brain Research, Seattle Children’s Research Institute; Lmx1aCre line^25^), Raj Awatramani (Department of Neurology and Center for Genetic Medicine, Northwestern University Feinberg Medical School Chicago, Illinois; β-catenin LOF mice^31^, Michael J. Holtzman (University of Washington, St. Louis), USA; Foxj1Cre line^36,37^. The Ai9 reporter mouse line was obtained from JAX labs Stock No. 007909. All animals were kept in an ambient temperature, with a 12hr light-dark cycle and food available ad libitum. Noon of the day of the vaginal plug was designated as embryonic day 0.5 (E0.5). Animals of both sexes were used for generating the control and Foxj1Cre::β-catenin GOF brain samples. For Lmx1aCre animals, since the transgene is on the X-Chromosome, females showed mosaic expression of the Cre recombinase due to random X inactivation. Cre-positive male embryos/pups were used for analysis except from Figure 6 and Supplementary Figure S7 where female embryos were analyzed. Controls were littermates wherever possible. Control embryos/pups carrying the Ai9 reporter to mark the Lmx1a or Foxj1 lineage carried the respective Cre transgenes. Breeding colonies were maintained with the female breeders carrying the Lmx1aCre and Foxj1Cre transgenes since this germline recombination occurs in the testes of these lines. Primers used for genotyping were: Cre F: 5’ATTTGCCTGCATTACCGGTC3’, Cre R: 5’ATCAACGTTTTCTTTTCGG3’, Cre positive DNA shows a band at 350bp. For detecting β-catenin LOF (Exon2-6 floxed) following primers were used: RM41 5’AAGGTAGAG TGATGAAAGTTGTT3’; RM42 5’CACCATGTCCTCTGTCTATTC3’; RM43 5’TACACTATTGAATCACAGGGACTT3’^31^. The PCR shows 221bp as Wild type band while 324bp as band corresponding to floxed allele. For genotyping β-catenin GOF (Exon 3 floxed) FP 5’ GCTGCGTGGACAATGGCTAC3’ and RP 5’GCTTTTCTGTCCGGCTCCAT3’^26^ was used which detects conditional allele at 550bp and wild type band at 350bp in PCR.

#### Human

Fetal brain specimens were obtained from either medically indicated or spontaneous abortions. The Institutional Ethical Review Board of the School of Medicine, University of Zagreb, and University Hospital Center Zagreb has approved the procedure for collecting postmortem brain samples (UHC Zagreb EP02/21AG; UZSM: 641-01/19-02/01). The fetal age was estimated based on crown-rump length (CRL, in mm), pregnancy records, and histological findings and expressed as GW. In addition, the correlation of maturational parameters (CRL, body mass, pregnancy records, and sonographic examination) revealed no evidence of growth retardation or malformations. Finally, specimens in which subsequent postmortem analysis revealed macroscopically or microscopically identified pathological changes, or postmortem autolytic changes, were excluded.

### Choroid plexus dissection

All dissections were performed in ice-cold PBS. For wholemount immunostained ChP preparations, telencephalic hemispheres were isolated using #5 forceps under a stereo zoom microscope (Nikon SMZ445). The rostral part of the ChP was gently extracted from the telencephalic ventricle using blunt forceps. The ChP was dissected using microdissection scissors while stabilizing the hemisphere using blunt forceps. For RNA Seq/ qPCR experiments, we used a line carrying the Ai9 reporter to avoid contamination from adjacent tissues. We examined the brains for Ai9 fluorescence before and after microdissection to ascertain that the Ai9-positive hem remained intact (Figure 2D). Furthermore, we isolated only the region of the ChP that was easily accessible and protruded into the telencephalic ventricle.

### Immunohistochemistry

#### Mouse sections and organoids

Mouse brain sections/organoid sections were mounted on plus slides (Catalogue number: EMS 71869-11) and dried for 2-3 hours. Slides were transferred to a slide mailer (Catalogue number: EMS 71549-08) containing PBS+0.01% TritonX-100 for 10 minutes followed by a wash with PBS+0.03% TritonX-100 for 5 minutes. For antigen retrieval, sections were boiled in a 10mM sodium citrate buffer (pH=6) at 90^0^C for 10 minutes using a water bath. Slides were cooled to room temperature and washed with PBS+0.01% TritonX-100 for 10 minutes. Blocking (5% horse serum/Lamb serum in PBS + 0.1% TritonX-100) for 1 hour followed by overnight primary antibody incubation at 4°C. Secondary antibody incubation was performed at room temperature for 2 hours followed by 3 washes with 1x PBS. Slides were mounted using vectashield mounting media (Vector laboratories H-1000-10) and imaged in an Olympus FluoView 1200 confocal microscope. The primary antibodies used were: LEF1 (rabbit, 1:200 CST catalogue # C12A5), β-CATENIN (mouse 1:200, BDbiosciences catalogue # 610153), β-CATENIN (rabbit 1:50, CST catalogue # 8814), RFP (rabbit, 1:200 Abcam catalogue # ab62341), FZD1 (rabbit, 1:100 catalogue # LS-A4150), AXIN2 (rabbit, 1:200 Abcam catalogue # ab185821), TTR (rabbit 1:100, DAKO catalogue # A0002), AQP1 (mouse, 1:200 SCBT catalogue # sc25287), OTX2(mouse, 1:200 SCBT catalogue # sc-514195), OTX2(1:500,invitrogen,PA5-39887), AQP1 (rabbit, 1:200,invitrogen catalogue # MA5-32593), E-CADHERIN (mouse, 1:200, SCBTcatalogue# sc21791), E-CADHERIN (mouse,1:100 BD Bio transductions, catalogue # 610182 0Cleaved CASPASE 3 (rabbit, 1:200 CST catalogue # 9664), Phospho HISTONE (rabbit, 1:200 CST catalogue # H0412), TBR2 (rabbit, 1:200 Abcam catalogue # ab23345), PAX6 (rabbit, 1:500 Abcam catalogue # ab195045), PROX1 (rabbit, 1:500 Millipore catalogue # ab5475), β-III TUBULIN (mouse, 1:100 Promega catalogue # G7128), BLBP (rabbit, 1:200 sigma catalogue # ABN14), ACTIN (mouse, 1:500 sigma catalogue # A2228), LAMININ (rabbit, 1:200 sigma catalogue # L9393). Secondary antibodies used were following: Goat Anti Rabbit Alexa fluor 488 (1:200, Invitrogen catalogue # A11034), Goat Anti mouse Alexa fluor 594 (1:200, Invitrogen catalogue # R37121), Goat Anti Rabbit Alexa fluor 568 (1:200, Invitrogen catalogue # A11011), Donkey Anti goat Alexa fluor 568 (1:200, Invitrogen catalogue # A32814), Donkey Anti rabbit Alexa fluor 647 (1:200, Invitrogen catalogue # A31573).

#### Human

The entire brains were fixed by immersion in 4% paraformaldehyde in 0.1 M phosphate buffer saline (pH=7.4). Subsequently, tissue blocks were embedded in paraffin wax. Serial sections were cut (5 μm) and stained. The cresyl violet staining was used to delineate cytoarchitectonic boundaries, while adjacent sections were labeled using immunofluorescence with various antibodies. The following primary antibodies were used: (1) rabbit polyclonal anti-Axin 2, (ab185821, Abcam); (2) the rabbit polyclonal Frizzled 1 antibody, FZD1 ( LS-A4150, LSBio); (3) the monoclonal rabbit LEF1 antibody (C12A5, CST); (4) the mouse monoclonal aquaporin1 antibody, AQP1 (sc-25287, SCBT); (5) Rabbit non-phospho (Active) βCatenin (D13A1, Cell Signaling, Leiden, Netherlands); (6) monoclonal mouse OTX2 antibody (MA5-15854, Invitrogen); (7) the mouse monoclonal Beta-Catenin, (M3539, Dako Agilent).The dewaxing of sections was performed in xylol for 5 min and a series of alcohol solutions (100%, 96%, 70%) for 2 min. Slides were then washed in PBS with 0.03% triton (10min). The antigen retrieval was performed in citrate buffer pH6.0 and microwave (800W) for 10 minutes. The slides were cooled at room temperature and immersed in a blocking solution (PBS containing 3 % bovine serum albumin, BSA, and 0.01 % Triton X-100, all from Sigma, St. Louis, MO) for 30 min. to prevent nonspecific background staining. Sections were then incubated with primary antibodies for 48 h at 4°C (AXIN2 1:100, FZD1 1:100, LEF1 1:100, AQP1 1:100, OTX2 1:50, β-Catenin cell signaling 1:400, β-Catenin Dako 1:200), washed and subsequently incubated with secondary anti-mouse or anti-rabbit antibodies diluted in PBS (at 1:1000) for 2 h at the room temperature. For visualization of the specific immune reactivity following secondary antibodies were used according to the laboratory protocol ^63^: the donkey anti-mouse IgG Alexa fluor-plus 488, donkey anti-rabbit IgG Alexa fluor-plus 647, donkey anti-rabbit IgG 488, donkey anti-mouse IgG Alexa-fluor 546 were used. Sections were covered with a mounting medium containing DAPI. Negative controls were included in all experiments by replacing the primary antibody with a blocking solution. No immunolabeling was detected in the control sections.

### In situ hybridization

Plasmids used to generate probes for *in situ* hybridization were kind gifts from Elizabeth Grove (*Axin2, Wnt3a, Ttr*), Cliff Ragsdale (*Fzd1, Fzd2a*) Kathleen Millen (*Lmx1a*). Fixed mouse brains were sectioned using a freezing microtome (Leica SM2000R Sliding Microtome) at a thickness of 20 μM. Sections were mounted on super frost plus slides (Catalogue #: EMS 71869-11), postfixed with 4% (w/v) paraformaldehyde for 15 minutes, washed three times with PBS. The sections were then treated with proteinase K dissolved in Tris-EDTA buffer (1 μg/ml). Post fixation was performed in 4% PFA for 15 minutes, and the sections were washed three times in PBS. Hybridization was performed for 16 h at 70°C in a buffer containing 50% (v/v) formamide, 2X SSC, and 1% (w/v) SDS. Digoxigenin (DIG)-labeled cRNA probes were used for hybridization and were prepared from the respective plasmids using the in-vitro transcription. After hybridization, three stringent washes (for 45 minutes each) were performed in solution X (50% formamide, 2X SSC, and 1% SDS) followed by a wash with 2XSSC and then 0.2XSSC. Blocking was performed for 1 hr with 10% horse serum in TBST (Tris-buffered saline pH 7.5 with 0.1% Tween-20). Sections were incubated in alkaline phosphatase-conjugated anti-DIG (Digoxygenin) antibody Fab fragments (1:5000; Roche, catalog #12486523) at 1:5000 in the blocking buffer and incubated at 4°C overnight. The color reaction was performed using NBT/BCIP substrate (Roche, 4-nitroblue tetrazolium chloride, catalog #70210625; 5-Bromo-4-chloro-3-indolyl phosphate, catalog #70251721). Counterstaining was performed using Fast Red (Sigma-Aldrich, catalog # N3020). Slides were dried and coverslipped using DPX mounting reagent (SDFCL, catalog #46029). Probes used for in-situ hybridization were generated using an in-vitro transcription reaction. Linearized templates for this process were generated by restriction digestion of plasmids (described in^64^). Plasmids used to generate probes for *in situ* hybridization were kind gifts from Elizabeth Grove (*Axin2, Wnt3a, Wnt2b, Ttr*), Cliff Ragsdale (*Fzd1, Fzd2a*) Kathleen Millen (*Lmx1a*).

### Image acquisition and analysis

Bright-field images were acquired using Zeiss Axioskop-2 plus microscope equipped with a Nikon DS-fi2 camera and associated software. Mouse sections were imaged in Olympus FluoView 1200 confocal microscope and the human brain sections were imaged in Olympus FluoView 3000 confocal microscope. All the organoid sections were imaged using the Andor Dragonfly spinning disk confocal microscope system. All the image analysis was done on Fiji-ImageJ, Imaris and/or Adobe Photoshop CS6.

For intensity quantification (Figure 1E), a line was drawn across the cell using the “line tool” option in Fiji (arrowheads mark the origin), and “grey values” were measured across the line.

For area measurements (Figure 3B) ROIs were drawn using the “freehand selection tool” around the *Ttr+* domains, and area was calculated. In the Lmx1a Cre:: β Catenin GOF brains where the *Ttr+* domains appear to be sparse, multiple ROIs were drawn separately and added in the ROI manager. The area for each *Ttr+* domain was calculated and summed to obtain the total area.

For nuclear intensity quantification of β-Catenin GOF (Figure 1J; Figure 2C and I; Supplementary Figure S2B and D; Figure 3D; Figure 5E) control and Lmx1a Cre::β Catenin GOF/LOF coronal sections were mounted on the same slide and processed for immunostaining. Images were acquired under similar PMT settings with a pixel distribution of 1024 X 1024 in “sequential scan mode”. The PMT setting was optimized such that saturation can be avoided. The z stack images are projected as “SUM slices’’and bit depth was set as “8bit”. From a composite image containing DAPI and markers, ROIs were drawn around the DAPI+ nuclei using the “freehand selection tool” in Fiji. When drawing the ROIs, the channel having markers was hidden using the “channel tool” to avoid any visual bias. The nucleus for which the boundary is not clear (overlapping with each other) was not scored. After completion of marking the ROIs using the DAPI channel, the marker channel was turned on and from this channel, “mean grey values” and “area” were measured using the “measure” option in the ROI manager. Our results are consistent with a recent study that examined the sub-cellular localization of β-Catenin and reported a substantial cytoplasmic component even when the nuclear accumulation is increased upon Wnt activation^65^.

For cell counts (Figure 3E; Figure 4C and E; Supplementary Figure S4B, C, and D; Figure 5D) “cell counter” plugin in Fiji was used. Cells for which the boundary was not adequately resolved were excluded. While scoring for DAPI+ or AI9+ cells, the channel containing the marker was hidden using the “channel tool” to avoid any visual bias. Similarly, while scoring markers, the Ai9 or DAPI channel was hidden using the “channel tool”. Similarly, ROI was drawn around the cytoplasm using the β-III TUB channel in Figure 6 and intensity was quantified from the Alexa 488 (TTR) and Alexa 647 (β-III TUBULIN) channels.

For Supplementary Figure 11C and D, a region of width 100µm was drawn using a “freehand line tool,” and the marker+ cells were scored using the “cell counter” plugin in Fiji.

Image stitching was performed for Figure 4D and F (low mags) using the “pairwise stitching” plugin in Fiji. The “subpixel accuracy” parameter was selected for all the stitching operations.

Nonlinear operations e.g. gamma adjustment were not performed in any of the figures. Brightness and contrast adjustments were performed identically for control and mutant conditions.

### Generation of hESC derived organoids

The human ES cell line WIBR3, obtained from the Whitehead Institute for Biomedical Research, (https://hpscreg.eu/cell-line/WIBRe001-A), was used in compliance with all applicable statutes and regulations and compliance with applicable guidelines (including the NAS Guidelines or guidelines of the International Society for Stem Cell Research as relevant to the conduct of human embryonic stem cell research [“ISSCR Guidelines”]. The line was cultured on an irradiated MEF (mouse embryonic fibroblast) feeder layer. To maintain the pluripotent state, a customized human Naïve media with 40 μM Rock Inhibitor Y-27632 was used as described in Karzbrun et al., 2018^66^. Differentiation of hES aggregates into Choroid plexus-like organoids was done using a protocol described in Sakaguchi et al. 2015^39^. Briefly, hESC colonies were grown in a Matrigel-coated plate till confluence. The cells were dissociated with trypsin and rapidly reaggregated in an ultra-low attachment 96 well V bottom plate (9000 cells per well) containing a media termed SA1 media. This media contains DMEM (Invitrogen cat # 11965092) supplemented with 20% (vol/vol) KSR (Invitrogen cat # 10828028), 0.1mM non-essential amino acids (Invitrogen cat # 11140050), 1mM pyruvate (Invitrogen cat # 11360070), 0.1mM 2-mercaptoethanol (Invitrogen cat # 21985023), 100U/ml penicillin and 100 mg/ml streptomycin. 0.3 mM IWR1e (tankyrase inhibitor, MedChemExpress cat# HY-12238) and 5 mM SB431542 (TGFβ inhibitor; MedChemExpress HY-10431) were added to culture from day 0 to day 18 (day of aggregation was defined as day 0). The culture media used after day 18 till day 40 was termed as SA2, which contains DMEM/F-12 (Invitrogen cat # 21331020), GlutaMAX(TM) (Gibco) supplemented with 1% N-2 supplement (Invitrogen cat # 17502048), 1% Chemically Defined Lipid Concentrate (Invitrogen cat # 11905031), fetal bovine serum (FBS; 10% vol/vol), 3 μM CHIR 99021 (GSK3 inhibitor; MedChemExpress cat # HY-10182) and 0.5 nM BMP4 (R&D cat # 314-BP-050) under normoxia condition. The medium was changed once every 2 days. From day 5 onwards, the plates were kept on an orbital shaker placed inside the incubator. On day 18 the organoids were transferred from 96 well plates to a 10 cm ultra-low attachment plate (6 organoids per plate) and kept on an orbital shaker inside the incubator. For sample collection, the organoids were rinsed with 1x PBS 2 times, then either flash-frozen for RNA-seq experiments or transferred to a small dish containing 4% PFA solution for immunohistochemistry.

### RNA sequencing, library preparation, and analysis of mouse tissues

The mouse ChP was dissected from E14.5 control and β-catenin GOF brains and stored in Trizol^Ⓡ^ reagent. Extracted RNA of 1 μg (RIN > 7.5, measured using the Agilent 2100 bioanalyzer) from ChPe dissected from 4 embryos was pooled for each of two biological replicates. After library preparation, sequencing was performed on the Illumina platform to achieve 100bp or 150bp reads to generate 30 Million paired-end reads per sample. FastQ QC was performed as described in ^67^, and reads > 30 Phred scores were aligned using HISAT2^68^. Feature counts were used to quantify the number of reads per transcript. Differential expression analysis was performed using DESeq2^69^ on the R platform (v3.6.3). Genes showing |log2 Fold change|≥1 were used for further analysis. Gene ontology analysis was performed using geneontology.org or gprofiler.org. Semantics were summarized using “REVIGO”^70^, and bar plots were created with GraphPad Prism V7.0 for Windows, GraphPad Software, San Diego California USA, www.graphpad.com). Heat maps and sample correlation plots were plotted using R studio. Gene-based heatmaps were plotted using normalized reads on Morpheus (Morpheus, https://software.broadinstitute.org/morpheus)

### RNA sequencing, library preparation, and analysis of organoids

Organoids generated in Figure 9 and 10 were pooled such that for one biological replicate, 3 organoids were used, and three such biological replicates were performed for day 18 and day 30 organoids. Bulk MARS-seq libraries were produced from 50ng of total RNA as previously described^71^. Libraries were then sequenced with 75 bp single-end read on Illumina Nextseq500 platform, and FASTQ files were processed and analyzed using UTAP^72^. Briefly, sequenced reads were trimmed using cutadapt (parameters: -a adaptor -a “A {10}” –times 2 -u 3 -u -3 -q 20 -m 25) and were mapped to hg38 indexed reference genome using STAR^73^ v2.4.2a with the following parameters: – alignEndsType EndToEnd, –out Filter Mismatch Nover Lmax 0.05, –two pass Mode Basic). UMIs were counted after marking duplicates using HTSeq-count in union mode. For the number of reads per gene, 1000bp of 3’ end Gencode annotated transcripts were counted. The total RNA-seq from 3 μM and 12 μM CHIR treated organoids (Figure 10) were done using TruSeq stranded mRNA library kit and sequenced by DNA Link Sequencing Lab, Korea, on NovaSeq 6000, 100PE. Data were analyzed with the UTAP pipeline^71^, where adapter trimming and reference genome alignment were performed as described above. Read count per gene was done with STAR. Normalization of the counts and differential expression analysis was performed using DESeq2^69^ with the parameters: betaPrior=True, cooks Cutoff=FALSE, independentFiltering=FALSE. Raw P values were adjusted for multiple testing using the procedure of Benjamini and Hochberg. Genes with log2FC >1 or <-1, p adjust < 0.05 and baseMean > 10 were considered as differentially expressed. GO Biological process overrepresentation test on differentially expressed genes between different time points of organoid growth was performed with the package clusterProfiler^74^.

### qPCR Analysis

#### Mouse

ChP samples from control and Lmx1aCre::β-catenin GOF E14.5 embryos were collected in a 1.5ml tube with 200 μl of Trizol reagent. ChP from 2 brains were pooled together and considered 1 biological replicate and 3 such biological replicates were performed. Total RNA was extracted using Trizol reagent following the manufacturer’s protocol. RNA concentration was measured using RNA-HS assay kit (catalog # Q32852) in QBIT-2 fluorometer (catalog # Q32866). cDNA was synthesized using SuperScript™ IV kit (catalog # 18091050). Real-Time qPCR reactions were performed in triplicates using KAPA SYBR FAST qPCR Kit (2X) (catalog # KK4601) on LightCycler^®^ 96 Real-time PCR system following the manufacturer’s recommendation. For each primer, the annealing temperature and primer concentration was optimized using gradient PCR. Melt curve analysis was performed to rule out the possibility of nonspecific amplification. GAPDH was used as a reference gene, and analysis was performed using the ΔΔ threshold cycle (Ct) method. The fold changes were represented as mean±SEM.

Primers: *Axin2*, FP: 5’-GAAGAAATTCCATACAGGAGGAT-3’; RP: 5’-GTCACTCGCCTTCTTGAAATAA-3; *Lef1*, FP: 5’-AGAAATGAGAGCGAATGTCGTAG-3’; RP: 5’-CTTTGCACGTTGGGAAGGA-3’; *Aqp1*, FP: 5’-CTGCTGGCGATTGACTACACTG-3’; RP: 5’-GGTTTGAGAAGTTGCGGGTGAG-3’; *Otx2*, FP: 5’-CAAAGTGAGACCTGCCAAAAAGA-3’; RP: 5’-TGGACAAGGGATCTGACAGTG-3’; *Slc12a2*, FP: 5’-GCAAGACTCCAACTCAGCCAC-3’; RP: 5’-ACCTCCATCATCAAAAAGCCACC-3’; *Slc12a4*, FP: 5’-GCCCCAACCTTACTGCTGAC-3’; RP: 5’-TCTCCTTTAGGCCGAGGGTG -3’; *Slc4a10*, FP: 5’-TTCAAGACCAGCCGCTATTT-3’; RP: 5’-GGATCCCAATGGCATAGTCA-3’; *Slc4a2*, FP: 5’-TCCAGAGCGAGCGGGTTATG-3’, RP: 5’-GAGGACTGGCGGTGGTACTCAAAGTC-3’; *Ttr*, FP: 5’-CCGTGTTAGCAGCTCAGGAA-3’; RP: 5’-GGGTTTTAGGAGCAGGGGAG-3’; *Gapdh*, FP: 5’-ATTCAACGGCACAGTCAAGG -3’; 5’-TGGATGCAGGGATGATGTTC-3’.

#### Organoids

Total RNA of Day 18 and Day 30 organoids was extracted using the RNeasy Mini kit (Qiagen, Germany) following the manufacturer’s protocol and followed by DNAse I treatment. RNA concentration was measured using Nanodrop (Thermo Scientific, MA, USA). cDNA was synthesized using M-MLV reverse transcriptase (M3682, Promega, Wisconsin, USA). Real-Time reactions were performed in triplicates using KAPA SYBR FAST qPCR Kit (2X) on QuantStudio 5 Real-time PCR system (Bio-Rad, CA, USA) following the manufacturer’s recommendation. Expression levels were normalized against GAPDH using the ΔΔ threshold cycle (Ct) method. The fold changes were represented as mean±SEM. Primers: *AQP1,* FP: 5’-TGGACACCTCCTGGCTATTG-3’; RP: 5’-GGGCCAGGATGAAGTCGTAG-3’; *LEF1,* FP: 5’-AGAACACCCCGATGACGGA-3’; RP: 5’-GAGGGTCCCTTGTTGTAGAGG-3’; *TTR,* FP: 5’ ATCCAAGTGTCCTCTGATGGT 3’; RP: 5’ GCCAAGTGCCTTCCAGTAAGA 3’; *OTX2,* FP: 5’-AGAGGACGACGTTCACTCG-3’; RP: 5’-TCGGGCAAGTTGATTTTCAGT-3’.

### Statistics

Biological replicates (N) denote samples obtained from individual brains. The total numbers of embryos/brains analyzed for immunohistochemistry or in situ hybridization for each genotype were:

69 Control brains (Figures 1-6)

10 Lmx1aCre::β-catenin LOF brains (Supplementary Figure S2 and 3)

46 Lmx1aCre::β-catenin GOF brains (Figure 2,3,4,6 and 7; Supplementary Figure S3,4,5,6 and 7)

7 Foxj1Cre::β-catenin GOF brains (Figure 5 and Supplementary Figure S5)

For qPCR and RNAseq, each biological replicate consisted of samples pooled from more than one brains that were processed as one sample. For qPCR (Figure 2 and 3), samples from 2 brains were pooled, while for RNA Seq (Figure 2 and 7, Supplementary Figure S8), samples from 4 brains were pooled and considered one biological replicate.

For organoids, one organoid was an independent biological replicate. A total of 12 control organoids (3 μM CHIR) were used for Figure 9. For Figure 10 and Supplementary Figure S10, 14 organoids (3 μM CHIR), 9 organoids (6 μM CHIR), 5 organoids (9 μM CHIR treated), and 14 organoids (12 μM CHIR treated), were used for immunohistochemistry analysis. For RNA-Seq experiments (Figures 9 & 10), 3 organoids (per treatment condition) are pooled as one biological replicate.

Detailed information about biological replicates and sample size was described in the corresponding figure legends. The genotypes could be distinguished by apparent phenotypic features, so it was not possible to perform the quantifications in a blinded manner. During image analysis, stringent measures were followed to avoid bias as described in the “Image analysis” section. All statistical analysis was performed in Graph Pad Prism V9, the exact test and “P-value” information were provided in corresponding figure legends. For all the graphs error bars indicate “standard error of mean”.

### Data availability

The RNA-seq data described in the manuscript are deposited in the GEO database (Accession number: GSE162784 and GSE162808). All other codes are available on request with the corresponding authors.

## Acknowledgments

We thank K. Millen (Seattle Children’s hospital) for the kind gift of the Lmx1aCre mouse line; Shital Suryavanshi and the animal house staff of the Tata Institute for Fundamental Research (TIFR) for excellent support. This work was supported by a Wellcome Trust-Department of Biotechnology India Alliance Early Career Fellowship (IA-E-12-1-500765; MC); by the Canada-Israel Health Research Initiative, jointly funded by the Canadian Institutes of Health Research, the Israel Science Foundation, the International Development Research Centre, Canada and the Azrieli Foundation (ST); intramural funds from TIFR-DAE (12-R&D-TFR-5.10-0100RTI2001).

Research in the Reiner lab is supported by ISF-National Science Foundation of China (NSFC) joint research program (Grant No. 2449/16), with the aid of a grant no. 2397/18 from the Canadian Institutes of Health Research (CIHR), the International Development Research Center (IDRC), the ISF, a grant from the Ministry of Science & Technology, Israel & The Ministry of Science and Technology of the People’s Republic of China., German-Israeli Foundation (GIF; Grant no. I-1476-203.13/2018), and United States-Israel Binational Science Foundation (BSF; Grant No. 2017006). In addition, by the Helen and Martin Kimmel Institute for Stem Cell Research, Nella and Leon Benoziyo Center for Neurological Diseases, David and Fela Shapell Family Center for Genetic Disorders Research, Brenden-Mann Women’s Innovation Impact Fund, Richard F. Goodman Yale/Weizmann Exchange Program, The Irving B. Harris Fund for New Directions in Brain Research, Irving Bieber, MD, and Toby Bieber, MD. Memorial Research Fund, Leff Family, Barbara & Roberto Kaminitz, Sergio & Sônia Lozinsky, Debbie Koren. Jack and Lenore Lowenthal, Dears Foundation. O. R. is the incumbent of the Berstein-Mason, professorial chair of Neurochemistry and Head of M. Judith Ruth Institute for Preclinical Brain Research.

The study on the human brains was financed by Croatian Science Foundation projects IP-2019-04-3182; DOK-2018-01-3771; DOK-2015-10-3939 to NJM and the Scientific Centre of Excellence for Basic, Clinical, and Translational Neuroscience (GA KK01.1.1.01.0007 funded by the European Union through the European Regional Development Fund). The collaboration was initiated throughout COST Action 16118.

We thank the members of Tole lab, Reiner lab, and Jovanov lab for their valuable inputs and discussion.

---

**Supplementary Figure S1:**
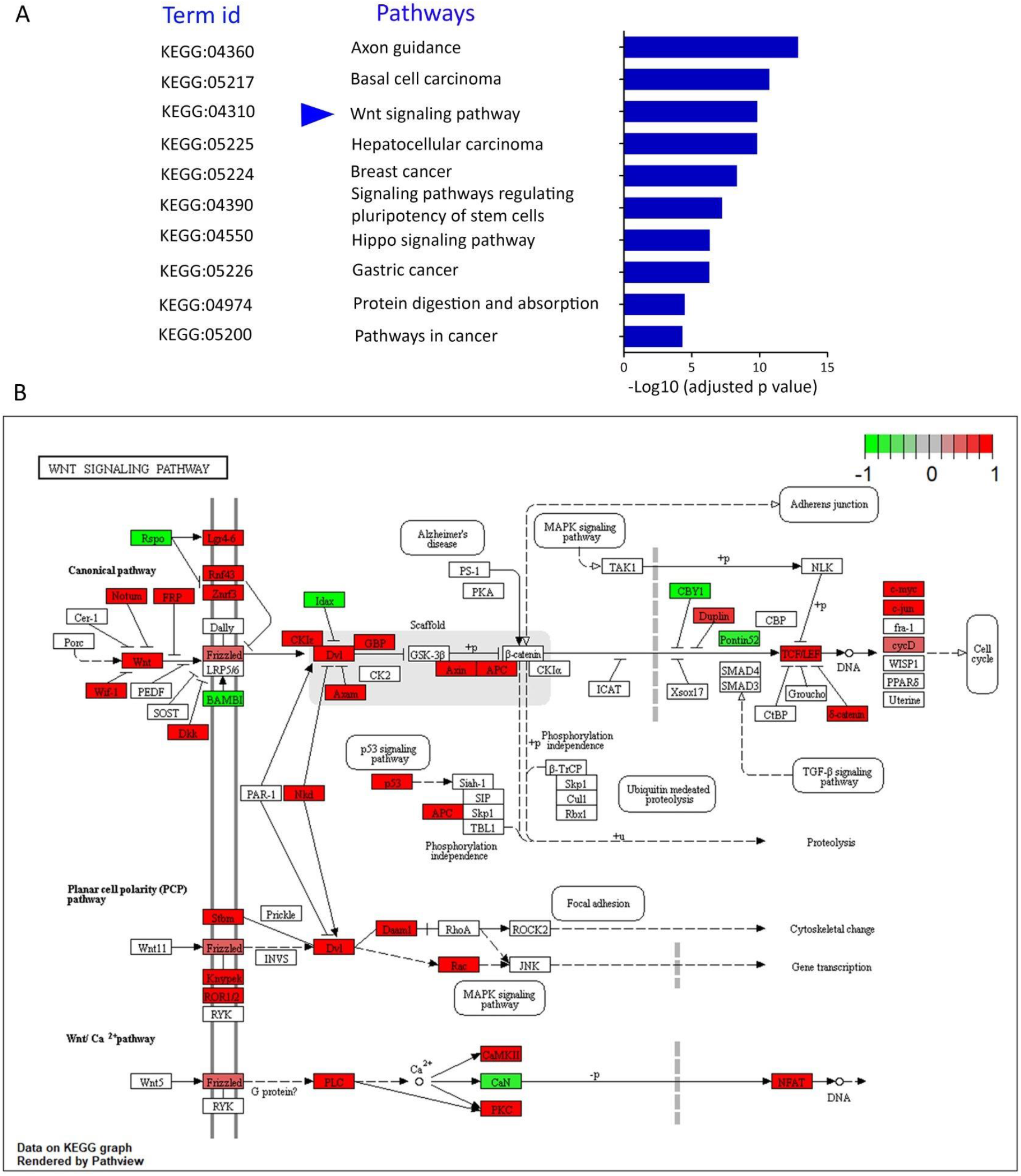
(A) KEGG pathway analysis identified the Wnt signaling pathway (blue arrowhead) among the top 10 dysregulated pathways. (B) Analysis of differentially expressed genes in the canonical Wnt signaling pathway and the planar cell polarity pathway in the control and β-CATENIN GOF ChP.

**Supplementary Figure S2.**
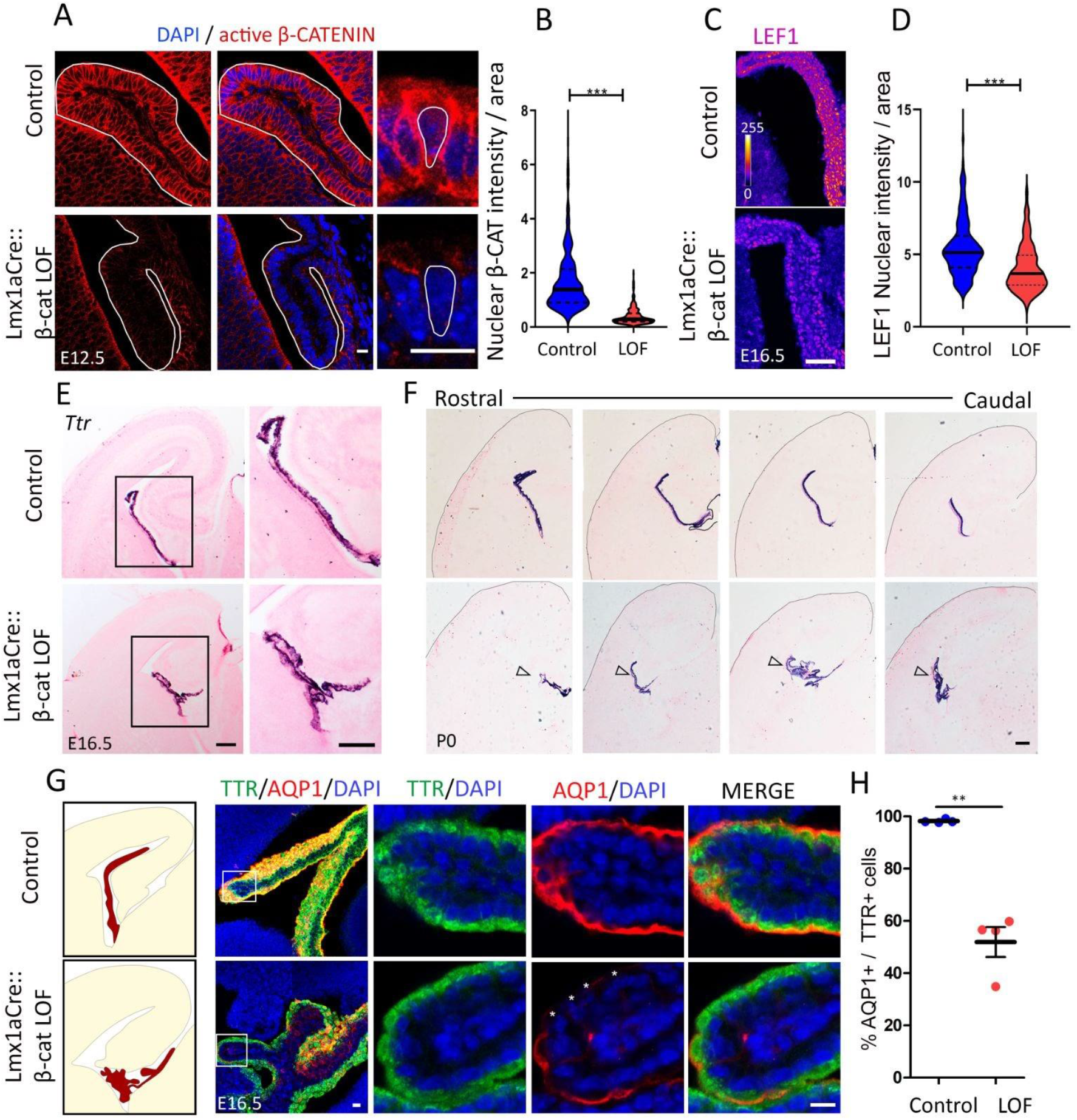
Loss of canonical Wnt signaling (β-CATENIN LOF) in the hem and ChPe disrupts choroid plexus development. (A) At E12.5, nuclear localization of active (non-phosphorylated) β-CATENIN was significantly reduced in the Lmx1aCre:: β-cat LOF ChPe compared with controls as seen by (A) immunohistochemistry and (B) a violin plot representing quantitation of 329 nuclei (control) and 302 nuclei (LOF); N=3 for each genotype. (C, D) At E16.5, LEF1 levels decrease (C), shown in a violin plot (D) representing quantitation of 401 nuclei (control) and 400 nuclei (LOF); N=3 for each genotype. (E, F) *Ttr* mRNA expression is maintained upon loss of β-CATENIN signaling at E16.5 and P0, even though there is substantial dysmorphia. (G, H) At E16.5, only 51.9 % of the TTR+ cells also display AQP1 (F, G). Scale bars: 10 μm (A and G); 50 μm (C); 100 μm (E and F).

**Supplementary Figure S3:**
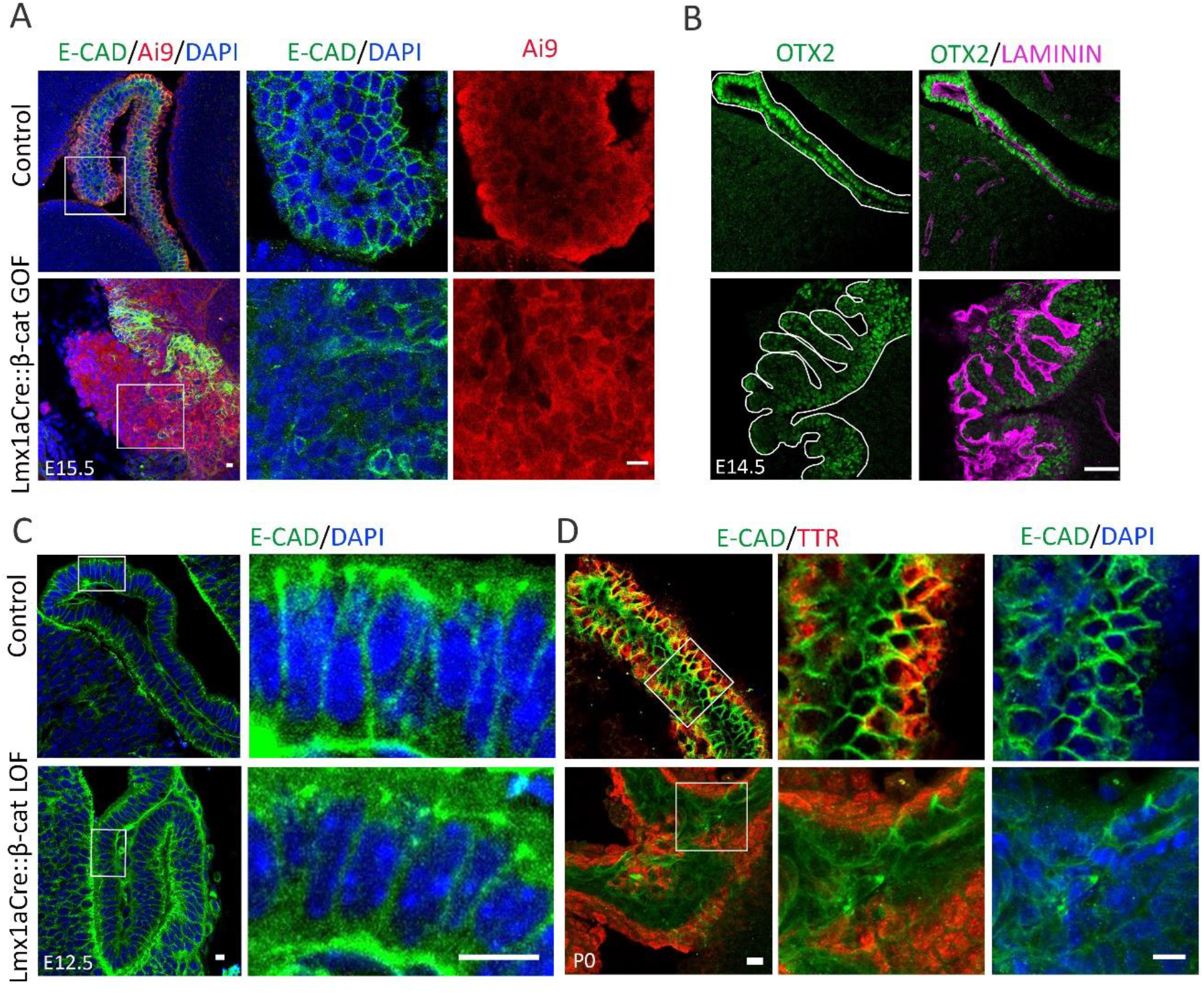
Cell Adhesion upon gain or loss of β-CATENIN. (A, B) At E14.5-E15.5, adherens junction marker E-CADHERIN (A) appears to be disorganized in the Lmx1aCre: β-cat GOF ChP compared with the control. Co-immunostaining for LAMININ and OTX2 (B) reveals a highly dysmorphic ChP with multiple folds and apparently reduced OTX2 in Lmx1aCre:: β-cat GOF embryos. (C) E-CADHERIN distribution appears similar at E12.5 in the Lmx1aCre: β-cat LOF ChP and control ChP. By birth (D) E-CADHERIN is disorganized in the β-cat LOF ChP which appears dysmorphic, although TTR labeling is maintained. Scale bars: 10 μm (A,C and D); 50 μm (B).

**Supplementary Figure S4:**
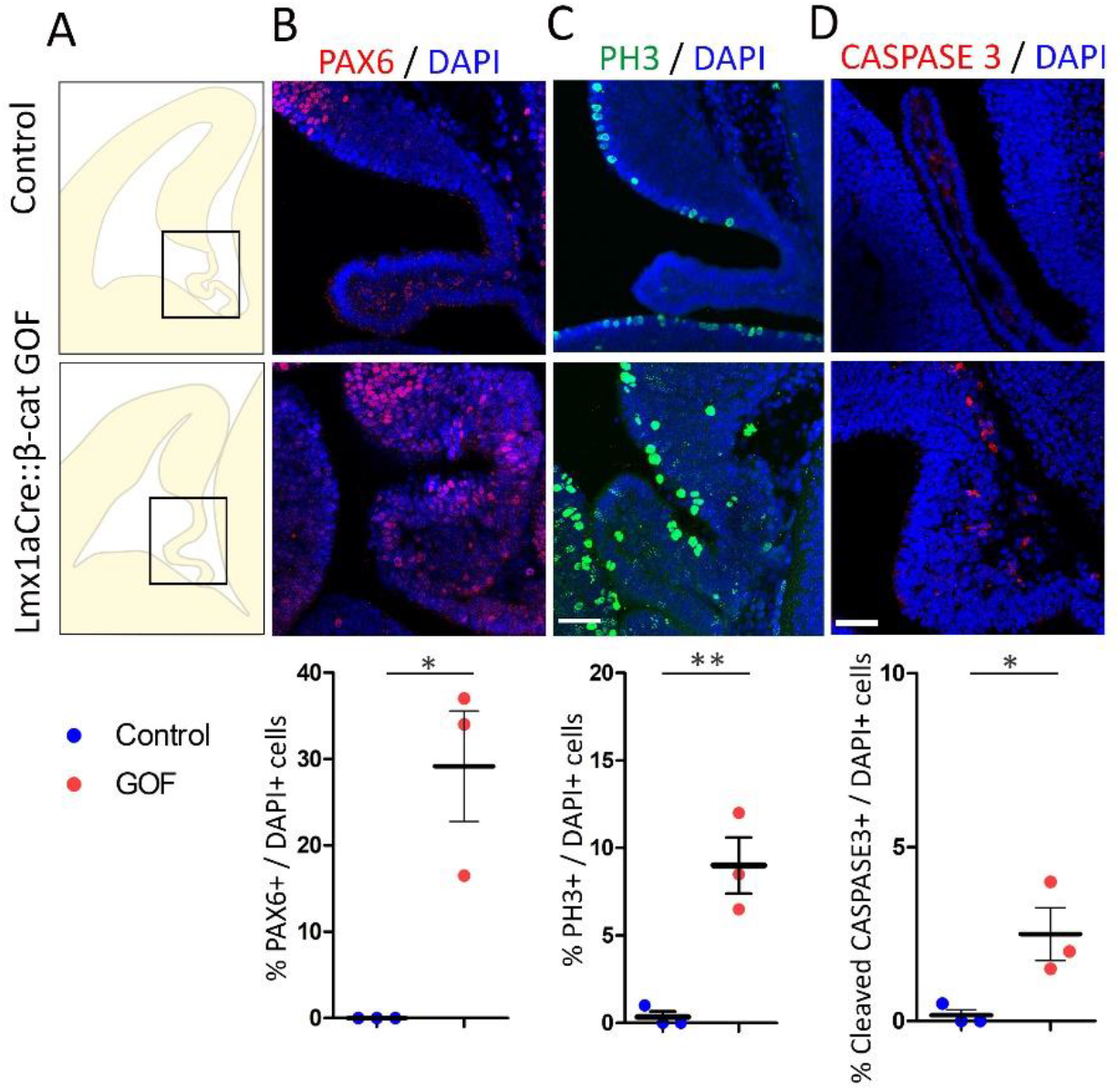
Proliferation and apoptosis in the choroid plexus upon stabilization of β-CATENIN. (A) Cartoons of E12.5/E13.5 mouse brain coronal sections showing the choroid plexus (Boxed region). (B) The neuronal apical progenitor marker PAX6 is present in 30% of the DAPI+ nuclei in the Lmx1aCre:: β-cat GOF ChP, whereas the control ChP does not display PAX6 labeling (N=3). (C) The cell proliferation marker PH3 labels 9% of the 422 DAPI+ cells scored in Lmx1aCre:: β-cat GOF ChP; while only 0.3% of the 400 DAPI+ cells are labeled in the control (N=3). (D) The apoptosis marker cleaved CASPASE 3 is detected in 2.5% of the 600 DAPI+ cells in Lmx1aCre:: β-cat GOF ChP and in 0.2% of 576 DAPI+ cells in the control (N=3). Statistical test: two-tailed unpaired Student’s t-test with unequal variance; * p < 0.05, ** p < 0.01, *** p < 0.001, ns if p-value> 0.05. Scale bars: 100 μm (C and D).

**Supplementary Figure S5:**
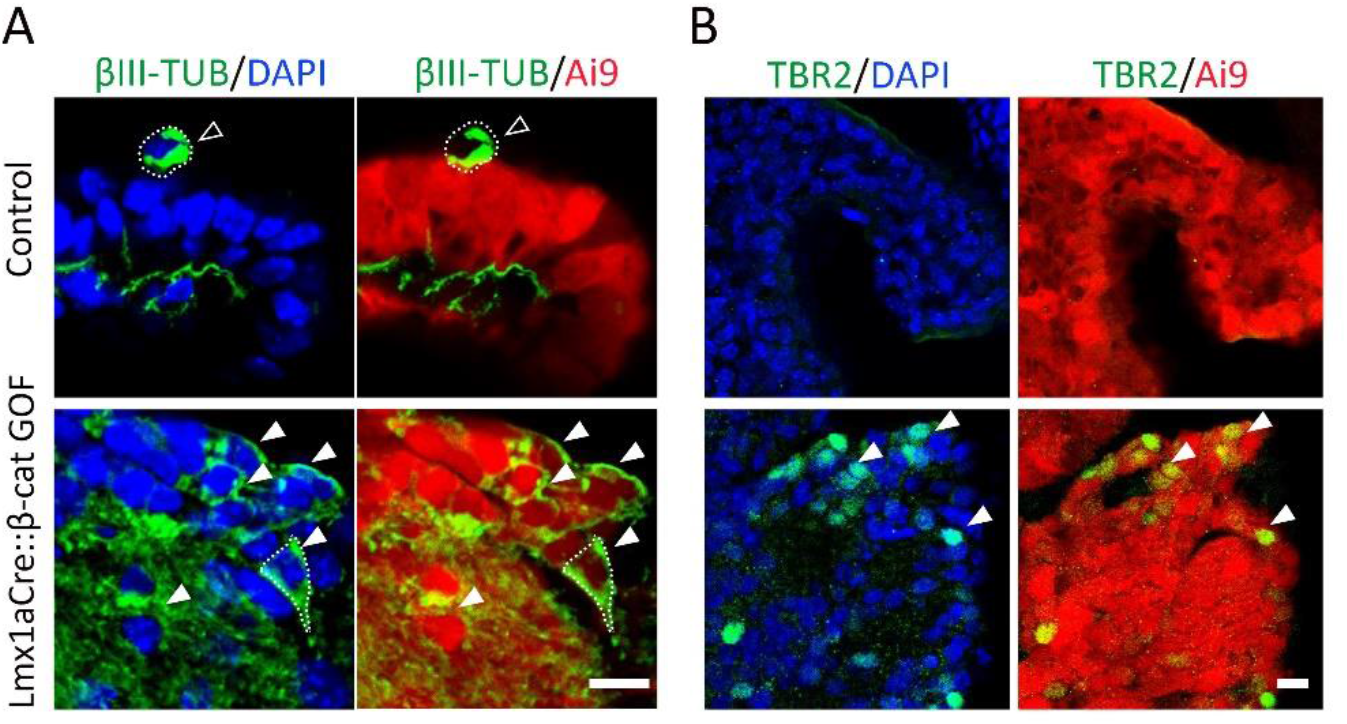
Neurons appear in the choroid plexus upon stabilization of β-CATENIN derived from the Lmx1a lineage. (A) At E16.5 a few βIII TUBULIN positive neurons are present in the control (Lmx1aCre::Ai9) ChP, but these do not co-localize with Ai9 and therefore do not stem from the Lmx1a lineage (open arrowheads, A). In contrast, multiple βIII TUBULIN-positive neurons and TBR2-positive cells are also Ai9-positive in Lmx1aCre::β-cat GOF:: Ai9 ChPe (arrowheads, A and B). Scale bars: 10 μm (A and B).

**Supplementary Figure S6:**
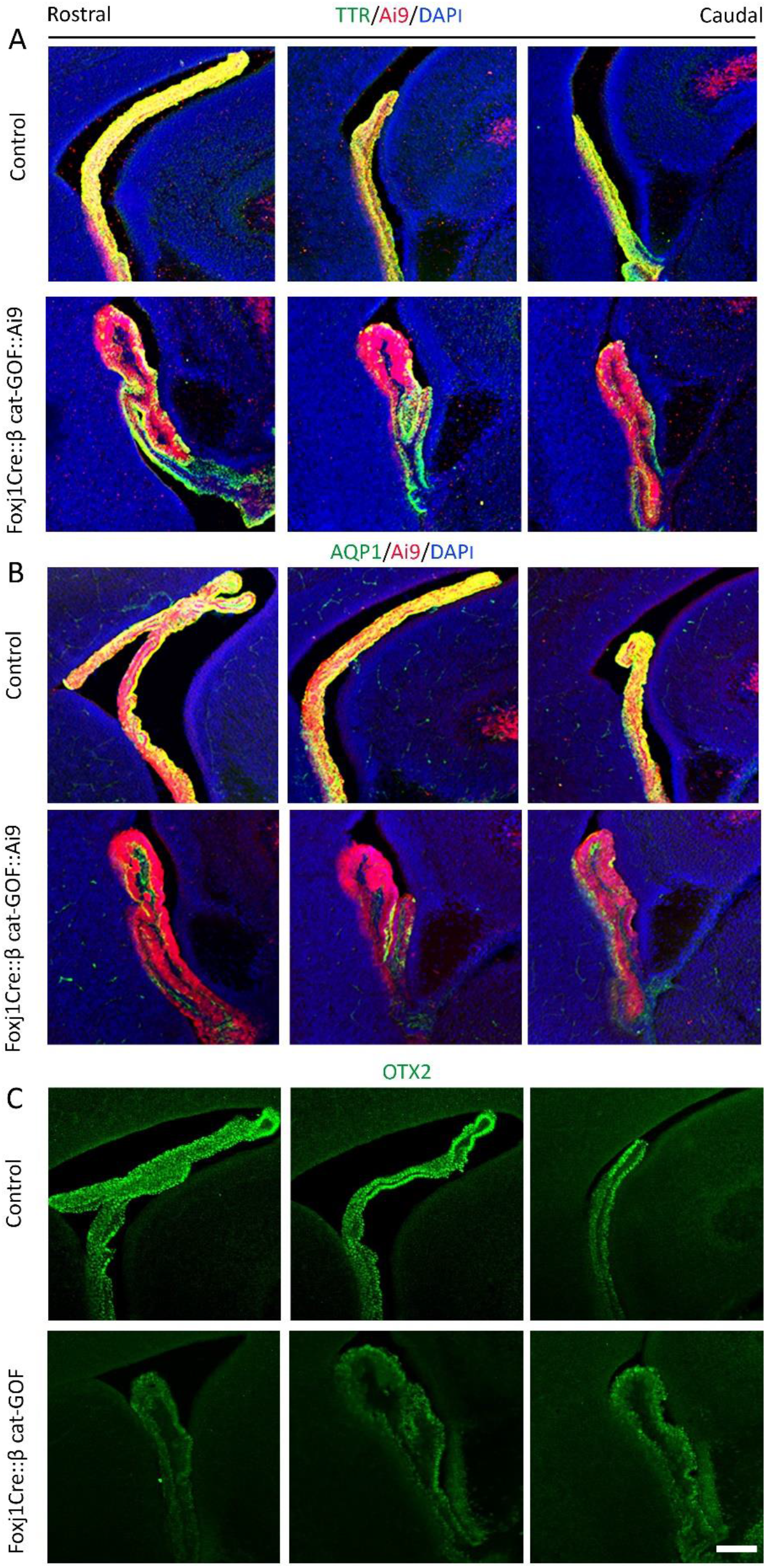
Rostrocaudal sections from E16.5 control and Foxj1Cre:: β-cat GOF embryos reveal TTR (A) and AQP1 (B) is lost from some regions of the ChP, whereas OTX2 (C) persists at lower levels in Foxj1Cre:: β-cat GOF brains. Scale bar: 100 μm.

**Supplementary Figure S7:**
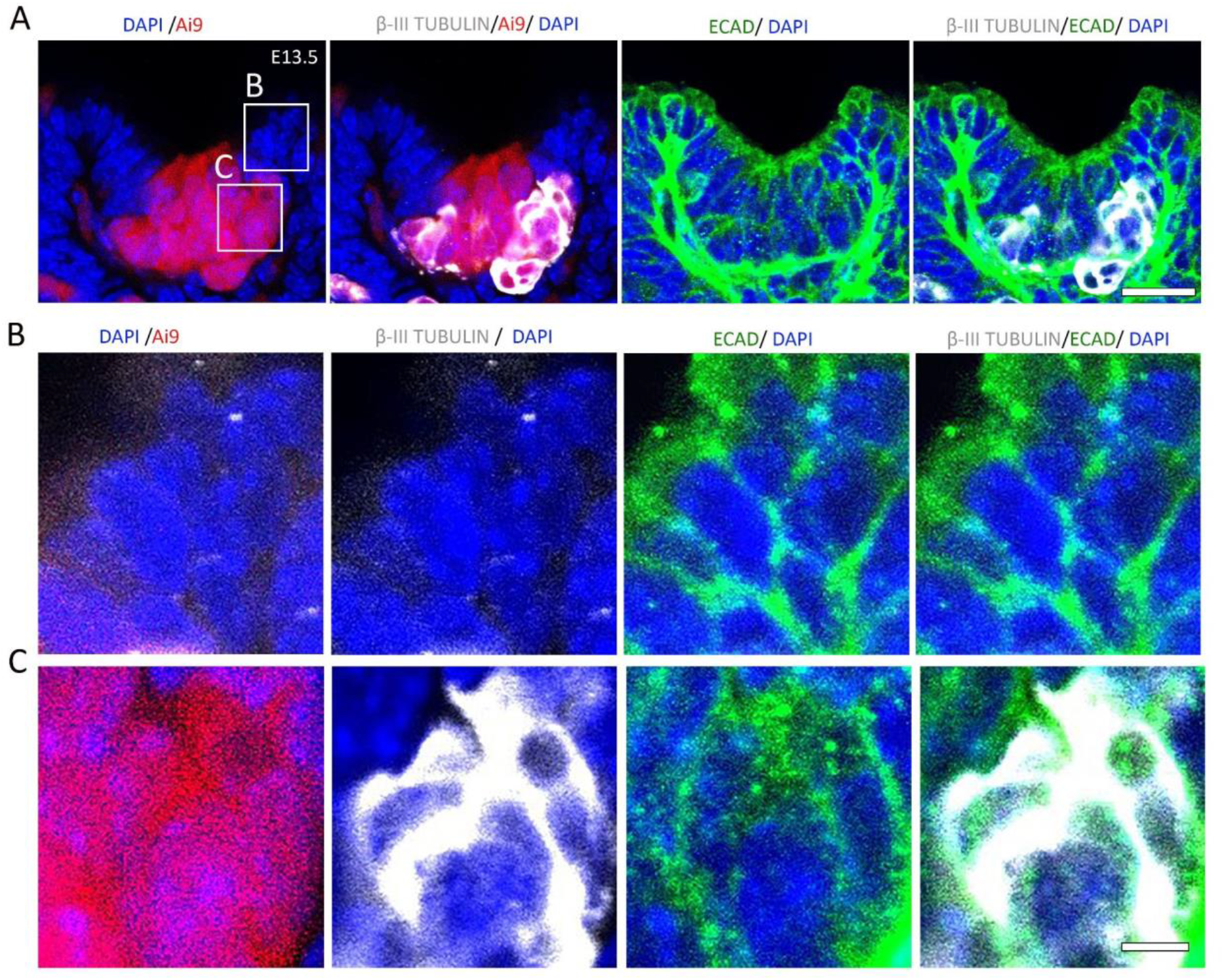
β-Catenin GOF causes upregulation of β-III TUBULIN in E-CADHERIN-positive ChPe cells. (A) Lmx1aCre::β-Catenin GOF::Ai9 female ChPe display mosaic Ai9+ patches interspersed with Ai9-patches. βIII TUBULIN appears selectively in the Ai9+ patches whereas E-CADHERIN staining is seen in both Ai9+ and Ai9-cells. (B) Ai9-internal control cells, do not display detectable βIII TUBULIN but contain E-CADHERIN. (C) Ai9+ cells display co-localization of β-III TUBULIN and E-CADHERIN. Scale bars: 5 μm (B and C); 20 μm (A).

**Supplementary Figure S8:**
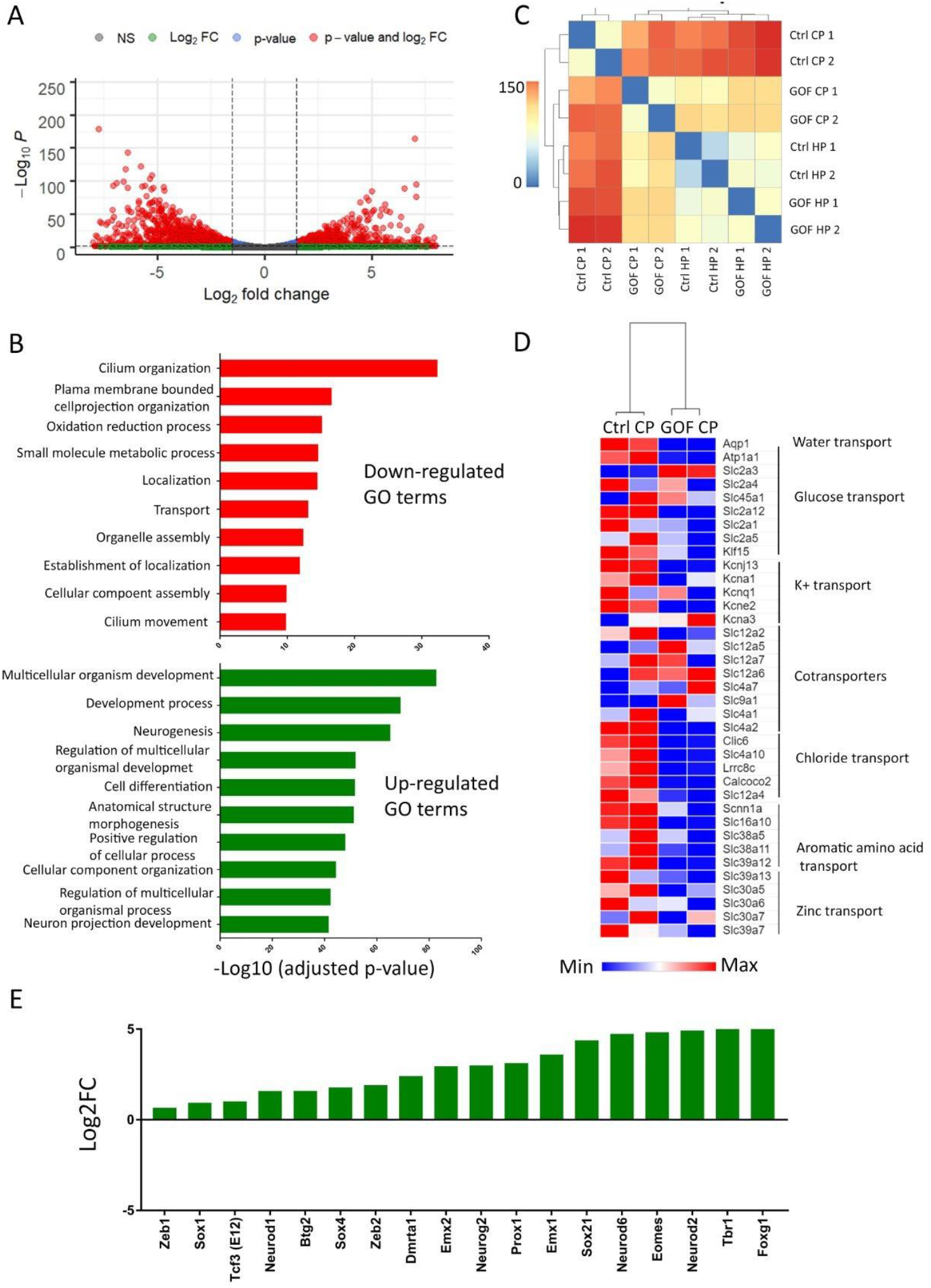
Analysis of differentially expressed gene sets in the control and Lmx1aCre:: β-catenin GOF ChP and Hippocampus. (A) Volcano plot and (B) GO analysis of differentially expressed genes in the control and β-catenin GOF ChP (C) Sample correlation matrix shows that the β-catenin GOF ChP is more similar to the control hippocampus (HP) than it is to the control ChP. (D) Heatmap of normalized read counts of a curated set of genes encoding different classes of transporters expressed in the normal ChP reveals they are uniformly downregulated in the β-catenin GOF ChP compared with the control. (E) Several pro-neuronal transcription factors belonging to the bHLH, Sox, and Zeb families are upregulated in Lmx1aCre::β-catenin GOF ChP.

**Supplementary Figure S9:**
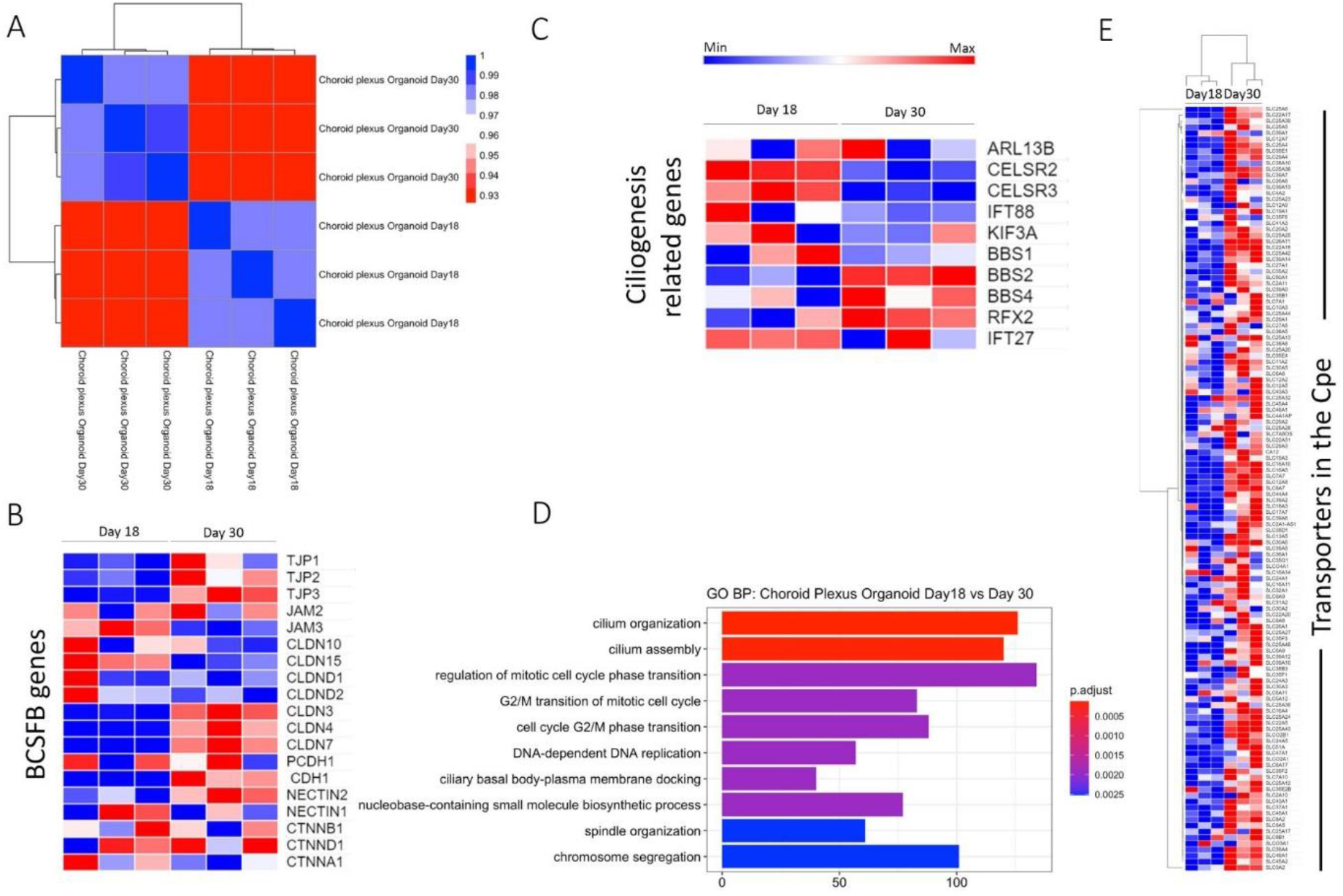
Characterization of ChPe-like differentiation in hESC organoids. (A-E) A comparison of transcriptomics data from day 18 and day 30 organoids. (A) Sample correlation matrix (B, C). Heat map of the normalized reads reveals that several BCSF genes are upregulated (B), and ciliogenesis-related genes are differentially regulated (C). (D) GO analysis (E) Heat map plotted from normalized reads shows an overall upregulation of transporter genes (belonging to the Slc gene family) in day 30 organoids.

**Supplementary Figure S10:**
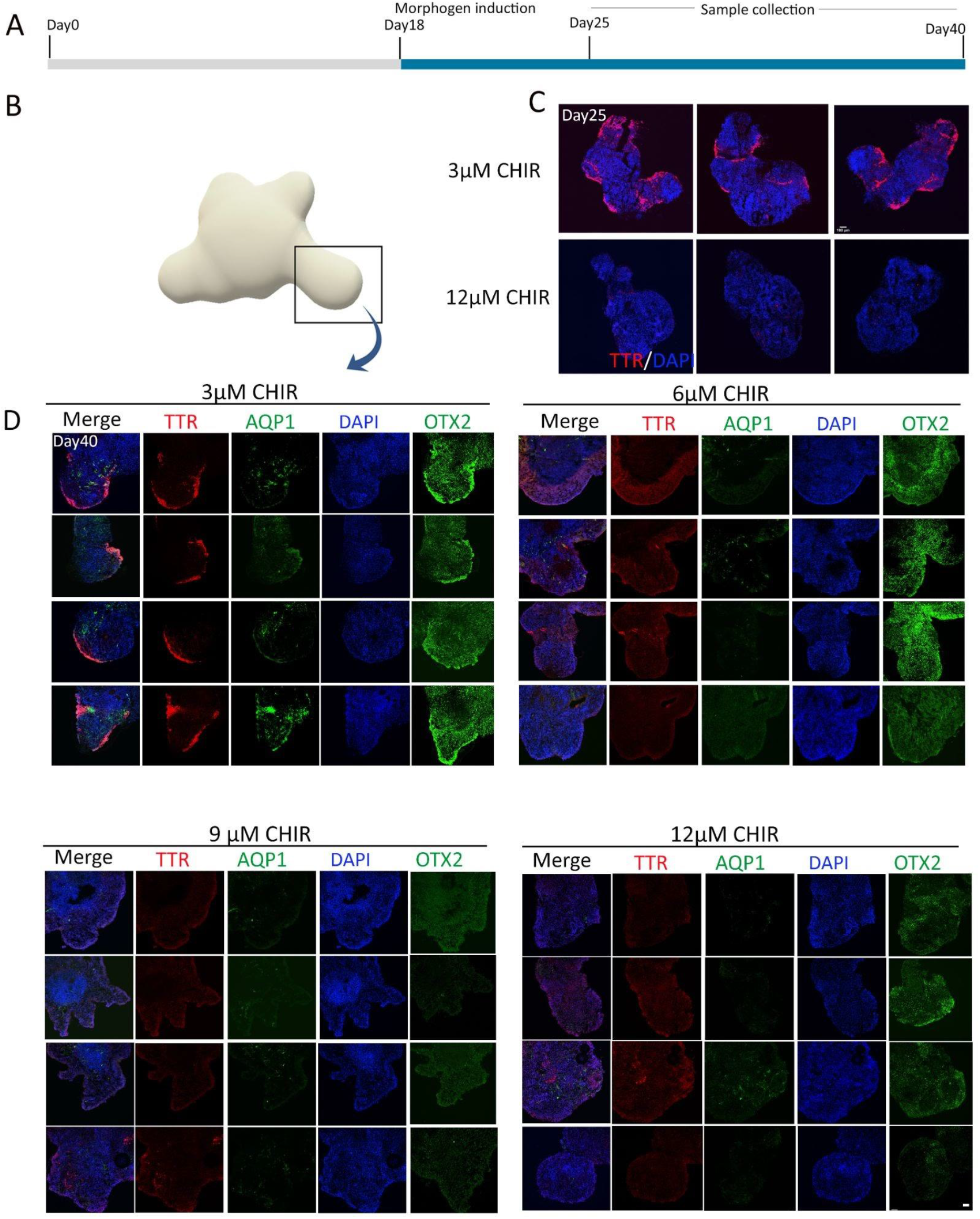
High activation of the canonical Wnt pathway in hESC-derived organoids causes suppression of ChPe markers. Additional examples of data shown in Figure 8 C and D. (A) Scheme showing the protocol for treatment with canonical Wnt agonist CHIR (C) ChP marker TTR is present in the periphery of day 25 organoids (N=3) exposed to 3 μM, but not 12 μM CHIR, from day 18. (D) High magnification images (boxed region shown in B) of serial sections of four examples of day 40 organoids exposed to 3 μM reveal TTR, AQP1, and OTX2 labeling in the periphery. This staining progressively decreases and is undetectable upon increasing exposure to higher concentrations of CHIR (6/ 9/ 12 μM). Scale bar: 100 μm.

**Supplementary Figure S11:**
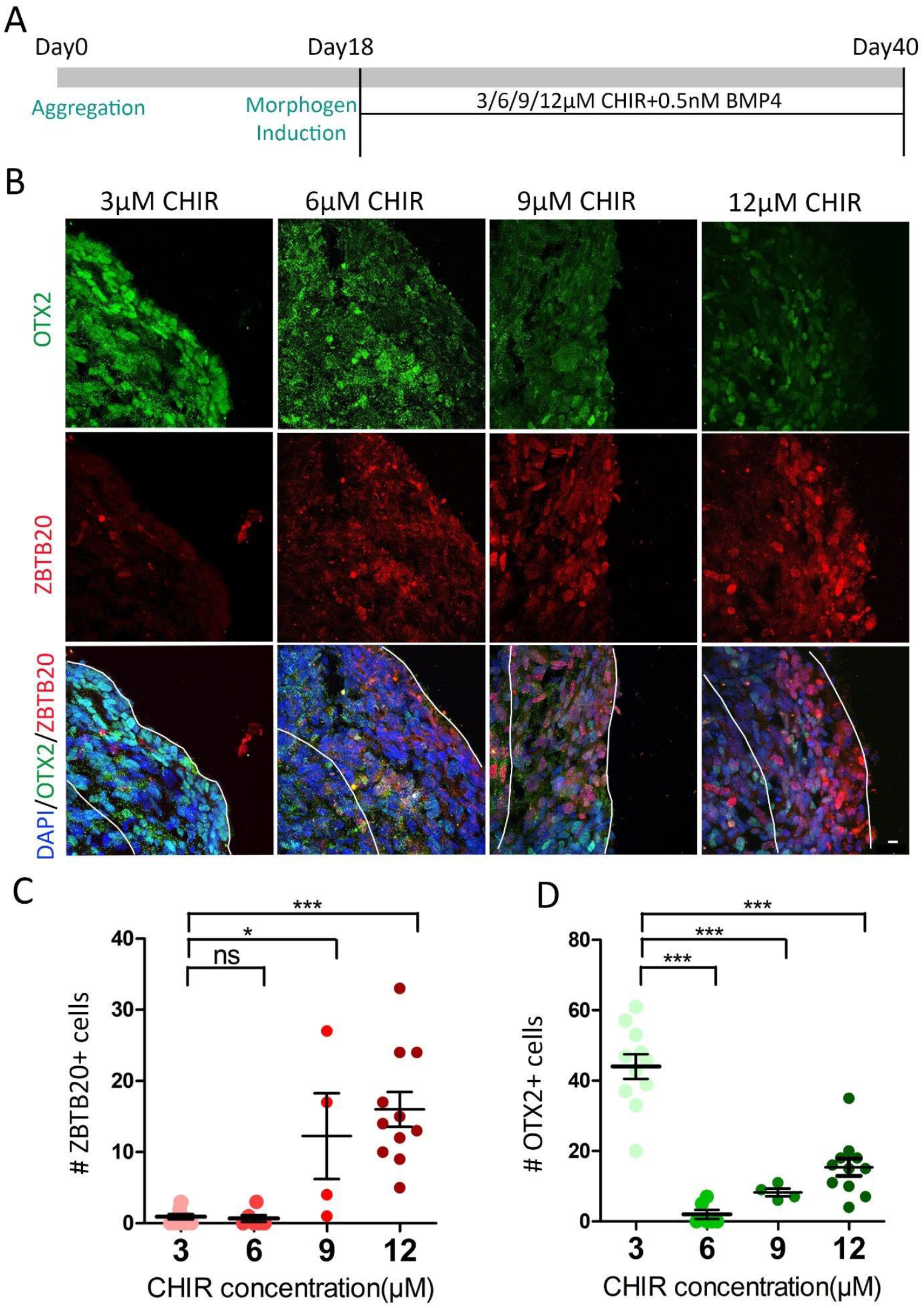
Hippocampus-specific marker ZBTB20 is upregulated, and OTX2 is downregulated upon high activation of the canonical Wnt pathway in hESC-derived organoids. (A) Schematic showing the organoid culture protocol and experimental design. (B) Immunofluorescence images of the periphery of an organoid display decreasing OTX2 and increasing ZBTB20 labeling upon exposure to increasing concentrations of CHIR. (C, D) Quantification of the number of ZBTB20+ and OTX2+ cells in organoids treated with 6/ 9/12 μM CHIR. N= 11 (3 and 12 μM CHIR); 6 (6 μM CHIR); 4 (9 μM CHIR). Statistical tests in C and D: two-tailed unpaired Student’s t-test with unequal variance, * p < 0.05, ** p < 0.01, *** p < 0.001, ns if p-value> 0.05. Scale bar: 10 μm.

## References

1. Johansson, P. A. The choroid plexuses and their impact on developmental neurogenesis. Front. Neurosci. 8, 1–9 (2014).

2. Lun, M. P., Monuki, E. S. & Lehtinen, M. K. Development and functions of the choroid plexus-cerebrospinal fluid system. Nat. Rev. Neurosci. 16, 445–457 (2015).

3. Zappaterra, M. W. & Lehtinen, M. K. The cerebrospinal fluid: Regulator of neurogenesis, behavior, and beyond. Cell. Mol. Life Sci. 69, 2863–2878 (2012).

4. Meeker, R. B., Williams, K., Killebrew, D. A. & Hudson, L. C. Cell trafficking through the choroid plexus. Cell Adhes. Migr. 6, 390–396 (2012).

5. Gu, X. et al. Inducible genetic lineage tracing of cortical hem derived Cajal-Retzius cells reveals novel properties. PLoS One 6, 1–9 (2011).

6. Louvi, A., Yoshida, M. & Grove, E. A. The derivatives of the Wnt3a lineage in the central nervous system. J. Comp. Neurol. 504, 550–569 (2007).

7. Yoshida, M., Assimacopoulos, S., Jones, K. R. & Grove, E. A. Massive loss of Cajal-Retzius cells does not disrupt neocortical layer order. Development 133, 537–545 (2006).

8. Roy, A., Gonzalez-Gomez, M., Pierani, A., Meyer, G. & Tole, S. Lhx2 regulates the development of the forebrain hem system. Cereb. Cortex 24, 1361–1372 (2014).

9. Liddelow, S. A., Dziegielewska, K. M., VandeBerg, J. L. & Saunders, N. R. Development of the lateral ventricular choroid plexus in a marsupial, Monodelphis domestica. Cerebrospinal Fluid Res. 7, 1–10 (2010).

10. Monuki, E. S., Porter, F. D. & Walsh, C. A. Patterning of the dorsal telencephalon and cerebral cortex by a roof plate-lhx2 pathway. Neuron 32, 591–604 (2001).

11. Konno, D. et al. The Mammalian DM Domain Transcription Factor Dmrta2 Is Required for Early Embryonic Development of the Cerebral Cortex. PLoS One 7, 1–13 (2012).

12. Grove, E. A., Tole, S., Limon, J., Yip, L. W. & Ragsdale, C. W. The hem of the embryonic cerebral cortex is defined by the expression of multiple Wnt genes and is compromised in Gli3-deficient mice. Development 125, 2315–2325 (1998).

13. Furuta, Y., Piston, D. W. & Hogan, B. L. M. Bone morphogenetic proteins (BMPs) as regulators of dorsal forebrain development. Development 124, 2203–2212 (1997).

14. Tole, S., Ragsdale, C. W. & Grove, E. A. Dorsoventral patterning of the telencephalon is disrupted in the mouse mutant extra-toes. Dev. Biol. 217, 254–265 (2000).

15. Hébert, J. M., Mishina, Y. & McConnell, S. K. BMP signaling is required locally to pattern the dorsal telencephalic midline. Neuron 35, 1029–1041 (2002).

16. Watanabe, M. et al. BMP4 sufficiency to induce choroid plexus epithelial fate from embryonic stem cell-derived neuroepithelial progenitors. J. Neurosci. 32, 15934–15945 (2012).

17. Eiraku, M. et al. Self-Organized Formation of Polarized Cortical Tissues from ESCs and Its Active Manipulation by Extrinsic Signals. Cell Stem Cell 3, 519–532 (2008).

18. Langford, M. B. et al. WNT5a Regulates Epithelial Morphogenesis in the Developing Choroid Plexus. Cereb. Cortex 30, 3617–3631 (2020).

19. Lee, S. M. K., Tole, S., Grove, E. & McMahon, A. P. A local Wnt-3a signal is required for development of the mammalian hippocampus. Development 127, 457–467 (2000).

20. Austinat, M. et al. Correlation between β-catenin mutations and expression of Wnt-signaling target genes in hepatocellular carcinoma. Mol. Cancer 7, 1–9 (2008).

21. Kim, S. & Jeong, S. Mutation hotspots in the β-catenin gene: Lessons from the human cancer genome databases. Mol. Cells 42, 8–16 (2019).

22. Li, L. et al. Sonic Hedgehog promotes proliferation of Notch-dependent monociliated choroid plexus tumour cells. Nat. Cell Biol. 18, 418–430 (2016).

23. Valenta, T. et al. Probing transcription-specific outputs of β-catenin in vivo. Genes Dev. 25, 2631–2643 (2011).

24. Chizhikov, V. V. & Millen, K. J. Control of roof plate formation by Lmx1a in the developing spinal cord. Development 131, 2693–2705 (2004).

25. Chizhikov, V. V. et al. Lmx1a regulates fates and location of cells originating from the cerebellar rhombic lip and telencephalic cortical hem. Proc. Natl. Acad. Sci. U. S. A. 107, 10725–10730 (2010).

26. Harada, N. et al. Intestinal polyposis in mice with a dominant stable mutation of the β-catenin gene. EMBO J. 18, 5931–5942 (1999).

27. Heiser, P. W., Lau, J., Taketo, M. M., Herrera, P. L. & Hebrok, M. Stabilization of β-catenin impacts pancreas growth. Development 133, 2023–2032 (2006).

28. Yan, D. et al. Elevated expression of axin2 and hnkd mRNA provides evidence that Wnt/β-catenin signaling is activated in human colon tumors. Proc. Natl. Acad. Sci. U. S. A. 98, 14973–14978 (2001).

29. Filali, M., Cheng, N., Abbott, D., Leontiev, V. & Engelhardt, J. F. Wnt-3A/β-catenin signaling induces transcription from the LEF-1 promoter. J. Biol. Chem. 277, 33398– 33410 (2002).

30. Hill, T. P., Später, D., Taketo, M. M., Birchmeier, W. & Hartmann, C. Canonical Wnt/β-catenin signaling prevents osteoblasts from differentiating into chondrocytes. Dev. Cell 8, 727–738 (2005).

31. Brault, V. et al. Inactivation of the β-catenin gene by Wnt1-Cre-mediated deletion results in dramatic brain malformation and failure of craniofacial development. Development 128, 1253–1264 (2001).

32. Caronia-Brown, G., Yoshida, M., Gulden, F., Assimacopoulos, S. & Grove, E. A. The cortical hem regulates the size and patterning of neocortex. Dev. 141, 2855–2865 (2014).

33. Mangale, V. S. et al. Lhx2 Selector Activity Specifies Cortical Identity and Suppresses Hippocampal Organizer Fate. Science (80-.). 319, 304 LP – 309 (2008).

34. Machon, O. et al. A dynamic gradient of Wnt signaling controls initiation of neurogenesis in the mammalian cortex and cellular specification in the hippocampus. Dev. Biol. 311, 223–237 (2007).

35. Dani, N. et al. A cellular and spatial map of the choroid plexus across brain ventricles and ages. Cell (2021) doi:10.1016/j.cell.2021.04.003.

36. Zhang, Y. et al. A transgenic FOXJ1-Cre system for gene inactivation in ciliated epithelial cells. Am. J. Respir. Cell Mol. Biol. 36, 515–519 (2007).

37. Jacquet, B. V. et al. Specification of a foxj1-dependent lineage in the forebrain is required for embryonic-to-postnatal transition of neurogenesis in the olfactory bulb. J. Neurosci. 31, 9368–9382 (2011).

38. Lun, M. P. et al. Spatially heterogeneous choroid plexus transcriptomes encode positional identity and contribute to regional CSF production. J. Neurosci. 35, 4903–4916 (2015).

39. Sakaguchi, H. et al. Generation of functional hippocampal neurons from self-organizing human embryonic stem cell-derived dorsomedial telencephalic tissue. Nat. Commun. 6, (2015).

40. Ying, Q. L. et al. The ground state of embryonic stem cell self-renewal. Nature 453, 519– 523 (2008).

41. Naujok, O., Lentes, J., Diekmann, U., Davenport, C. & Lenzen, S. Cytotoxicity and activation of the Wnt/beta-catenin pathway in mouse embryonic stem cells treated with four GSK3 inhibitors. BMC Res. Notes 7, 1–8 (2014).

42. Xie, Z. et al. Zbtb20 is essential for the specification of CA1 field identity in the developing hippocampus. Proc. Natl. Acad. Sci. U. S. A. 107, 6510–6515 (2010).

43. Galceran, J., Miyashita-Lin, E. M., Devaney, E., Rubenstein, J. L. R. & Grosschedl, R. Hippocampus development and generation of dentate gyrus granule cells is regulated by LEF1. Development 127, 469–482 (2000).

44. Ivaniutsin, U., Chen, Y., Mason, J. O., Price, D. J. & Pratt, T. Adenomatous polyposis coli is required for early events in the normal growth and differentiation of the developing cerebral cortex. Neural Dev. 4, (2009).

45. Oosterwegel, M. et al. Differential expression of the HMG box factors TCF-1 and LEF-1 during murine embryogenesis. Development 118, 439–448 (1993).

46. Fischer, T., Guimera, J., Wurst, W. & Prakash, N. Distinct but redundant expression of the Frizzled Wnt receptor genes at signaling centers of the developing mouse brain. Neuroscience 147, 693–711 (2007).

47. Kaiser, K., Jang, A., Lun, M. P., Procházka, J. & Machon, O. MEIS-WNT5A axis regulates development of 4. (2020).

48. Huelsken, J., Vogel, R., Erdmann, B., Cotsarelis, G. & Birchmeier, W. β-Catenin controls hair follicle morphogenesis and stem cell differentiation in the skin. Cell 105, 533–545 (2001).

49. Kajino, Y. et al. β-Catenin gene mutation in human hair follicle-related tumors. Pathol. Int. 51, 543–548 (2001).

50. Kemler, R. et al. Stabilization of β-catenin in the mouse zygote leads to premature epithelial-mesenchymal transition in the epiblast. Development 131, 5817–5824 (2004).

51. Wang, Y. et al. Beta-catenin signaling regulates barrier-specific gene expression in circumventricular organ and ocular vasculatures. Elife 8, 1–36 (2019).

52. Li, C. et al. Apc deficiency alters pulmonary epithelial cell fate and inhibits Nkx2.1 via triggering TGF-beta signaling. Dev. Biol. 378, 13–24 (2013).

53. Li, J. et al. Integrative genomic analysis of early neurogenesis reveals a temporal genetic program for differentiation and specification of preplate and Cajal-Retzius neurons. PLoS Genetics vol. 17 (2021).

54. Imayoshi, I., Shimogori, T., Ohtsuka, T. & Kageyama, R. Hes genes and neurogenin regulate non-neural versus neural fate specification in the dorsal telencephalic midline. Development 135, 2531–2541 (2008).

55. Ernst, S. A., Palacios, J. R. & Siegel, G. J. Immunocytochemical localization of Na+,K+-ATPase catalytic polypeptide in mouse choroid plexus. J. Histochem. Cytochem. 34, 189–195 (1986).

56. Amin, M. S., Reza, E., Wang, H. & Leenen, F. H. H. Sodium transport in the choroid plexus and salt-sensitive hypertension. Hypertension 54, 860–867 (2009).

57. Cornejo, I. et al. Tissue distribution of Kir7.1 inwardly rectifying K+ channel probed in a knock-in mouse expressing a haemagglutinin-tagged protein. Front. Physiol. 9, 1–12 (2018).

58. Speake, T., Kibble, J. D. & Brown, P. D. Kv1.1 and Kv1.3 channels contribute to the delayed-rectifying K + conductance in rat choroid plexus epithelial cells. Am. J. Physiol. - Cell Physiol. 286, 611–620 (2004).

59. Steffensen, A. B. et al. Cotransporter-mediated water transport underlying cerebrospinal fluid formation. Nat. Commun. 9, 1–13 (2018).

60. Kim, J. & Jung, Y. Increased aquaporin-1 and Na+-K+-2Cl-cotransporter 1 expression in choroid plexus leads to blood-cerebrospinal fluid barrier disruption and necrosis of hippocampal CA1 cells in acute rat models of hyponatremia. J. Neurosci. Res. 90, 1437–1444 (2012).

61. Björklund, P., Lindberg, D., Åkerström, G. & Westin, G. Stabilizing mutation of CTNNB1/beta-catenin and protein accumulation analyzed in a large series of parathyroid tumors of Swedish patients. Mol. Cancer 7, 1–8 (2008).

62. Pellegrini, L. et al. Human CNS barrier-forming organoids with cerebrospinal fluid production. Science (80-.). 369, 1–21 (2020).

63. Culjat, M. & Milošević, N. J. Callosal septa express guidance cues and are paramedian guideposts for human corpus callosum development. J. Anat. 235, 670–686 (2019).

64. Tole, S. & Patterson, P. H. Regionalization of the developing forebrain: A comparison of FORSE-1, Dlx-2, and BF-1. J. Neurosci. 15, 970–980 (1995).

65. de Man, S. M. A., Zwanenburg, G., van der Wal, T., Hink, M. A. & van Amerongen, R. Quantitative live-cell imaging and computational modelling shed new light on endogenous wnt/ctnnb1 signaling dynamics. Elife 10, (2021).

66. Karzbrun, E., Kshirsagar, A., Cohen, S. R., Hanna, J. H. & Reiner, O. Human brain organoids on a chip reveal the physics of folding. Nature Physics vol. 14 515–522 (2018).

67. Anders, S., Pyl, P. T. & Huber, W. HTSeq-A Python framework to work with high-throughput sequencing data. Bioinformatics 31, 166–169 (2015).

68. Kim, D., Langmead, B. & Salzberg, S. L. HISAT: A fast spliced aligner with low memory requirements. Nat. Methods 12, 357–360 (2015).

69. Love, M. I., Huber, W. & Anders, S. Moderated estimation of fold change and dispersion for RNA-seq data with DESeq2. Genome Biol. 15, 1–21 (2014).

70. Supek, F., Bošnjak, M., Škunca, N. & Šmuc, T. Revigo summarizes and visualizes long lists of gene ontology terms. PLoS One 6, (2011).

71. Jaitin, D. A. et al. Massively Parallel Single-Cell RNA-Seq for Marker-Free Decomposition of Tissues into Cell Types. Science (80-.). 343, 776 LP – 779 (2014).

72. Kohen, R. et al. UTAP: User-friendly Transcriptome Analysis Pipeline. BMC Bioinformatics 20, 1–7 (2019).

73. Dobin, A. et al. STAR: Ultrafast universal RNA-seq aligner. Bioinformatics 29, 15–21 (2013).

74. Yu, G., Wang, L. G., Han, Y. & He, Q. Y. ClusterProfiler: An R package for comparing biological themes among gene clusters. Omi. A J. Integr. Biol. 16, 284–287 (2012).

